# Stroke and Motor Recovery are Associated with Regional and Age-Specific Changes in Periodic and Aperiodic Cortical Activity

**DOI:** 10.1101/2024.11.07.622359

**Authors:** Asher J. Albertson, Eric. C. Landsness, Margaret Eisfelder, Brittany M Young, Bradley Judge, Matthew R. Brier, Matthew J. Euler, Steven C. Cramer, Jin-Moo Lee, Keith R Lohse

## Abstract

Normal aging is associated with widespread changes in neuronal structure, function, and activity. The consequences of focal brain injury on global neuronal activity in aged individuals are poorly understood. Historically, stroke and aging have been associated with changes in narrow-band periodic neuronal activity, however recent work has highlighted the importance of broad-band aperiodic activity. Aperiodic activity is represented by the 1/f slope of power spectral density generated by cortical activity.

Abnormalities in aperiodic activity have been identified in psychiatric disorders and stroke, are associated with cognitive dysfunction in aging, and have been hypothesized to reflect changes in excitation/inhibition balance. Here we sought to further explore changes in both periodic and aperiodic cortical activity in neurotypical intact young and aged healthy individuals and individuals with stroke. We compared “resting state” electroencephalograms from all participants after applying the *specparam* algorithm, which decomposes the power spectrum into aperiodic and periodic components. We also correlated stroke outcomes using previously obtained tests of motor outcome (box and block) to average whole cortex spectral slopes within the stroke group. Consistent with prior work we found a significant flattening (decrease in exponent) of power spectral slope with normal aging. We also found that both aging and stroke were associated with fewer periodic peaks within the power spectrum. Interestingly, we found that stroke was associated with a significant increase in spectral slope, but age moderated this effect. Younger stroke patients showed minimal difference in slope while older stroke patients had significantly steeper slopes (opposite to the direction in normal aging). Using MRIs from stroke participants we investigated the lesion locations most associated with changes in slope. Interestingly deep lesions were observed to have the greatest influence on cortical spectral slope. Finally, slope in the stroke group was correlated with performance on a test of manual dexterity, however this correlation was much more significant in aged individuals. Our data suggest that stroke in the aged brain has unique effects on aperiodic activity possibly reflecting unique influence of injury on cerebral excitation/inhibition balance in aged individuals and that the degree of these changes may be related to stroke outcomes.

## 1. Introduction

The impact of stroke is particularly significant in older individuals, among whom the incidence is substantially greater when compared to younger individuals^1^. Age is the largest non-modifiable risk factor for stroke^2-5^. Ischemic stroke survival is markedly reduced in aged individuals, and long-term disability is significantly increased^6^. Ischemic stroke is also associated with an increased risk of dementia^7,8^, with age being a significant risk factor for combined death and dementia^9^ after stroke. In contrast, older age is associated with lower risks of both post-stroke epilepsy^10^ and depression^11^. The neurologic and physiologic drivers of age-related differences in post-stroke cerebral function, and thus outcomes, are poorly understood.

Normal aging is associated with substantial changes in neuronal and cerebral physiology. Brain weight^12^, myelinated fibers^13^, and cortical thickness^14^ all decline with age. These anatomical changes are accompanied by changes in synaptic dynamics^15^, neuron structure^16^, synaptic plasticity^17,18^, and network functional connectivity^19-21^. Numerous clinical and preclinical studies have demonstrated that stroke also causes short- and long-term changes in brain structure^22,23^, synaptic function^24-26^, and network connectivity^27-29^. The differential effect of stroke on brain structure and function in aged individuals is less well understood.

Age-related changes in neuronal structure and function are reflected in changes in neuronal activity. Periodic spontaneous neuronal activity (defined as rhythmic neuronal oscillations within canonical frequency bands) changes substantially with age, and these changes may be indicators of cognitive performance^30^. Recent work from healthy young and aged adults showed that aging is associated with a relatively linear decline in slow frequency (0.5-6.5Hz) cortical activity. Interestingly, increased low frequency power in older but not younger adults was associated with better cognitive performance^31^.

This was speculated to reflect global changes in synaptic function^31^. Stroke also causes significant changes in spontaneous neuronal activity. Hyper-acute stroke is associated with broadband decreases in cortical spectral power, mostly in higher frequencies^32^. Dynamic increases in low frequency power have been reported at various time points after stroke^33-35^. Interestingly, increased low frequency activity after stroke is associated with both larger injury size and greater motor recovery^36^. Whether stroke uniquely affects spontaneous periodic activity in aged vs. young individuals is incompletely characterized.

While changes in periodic cortical activity in aging and stroke have received significant attention, a separate body of work has demonstrated the independent importance of aperiodic activity in cerebral function^37-40^. Aperiodic activity is represented by the 1/*f* slope of power spectral density generated by cortical activity. Cortical field potentials measured by electroencephalography (EEG) exhibit a characteristic negative slope, with the greatest power at lower frequencies^40^. Disruptions in 1/f spectral slope have been identified in neuropsychiatric disorders including schizophrenia^41^ and ADHD^42^.

Simulation and *in vivo* data have suggested that 1/*f* slope inversely reflects excitation/inhibition balance^43^. Similar to periodic activity, there are significant changes in aperiodic activity with age, and these changes correlate to performance on cognitive tests ^39,44^. Recent work has demonstrated that spectral slowing after stroke is a result of both periodic (diminished high frequency oscillations) and aperiodic (increased spectral slope)^45^ changes.

Given the relevance of neuronal activity^46^, synaptic function^47^, and, importantly, disrupted excitation/inhibition balance^48^, to ischemic brain injury and recovery, we sought to further investigate changes in post-stroke cortical activity across both the periodic and aperiodic components in aged and young individuals. We compared resting state EEGs from both healthy young and aged individuals to young and aged individuals with stroke. We applied the *specparam* algorithm (formerly *fitting oscillations with one-over-f*, or *FOOOF*) to each EEG timeseries at each electrode to quantify the aperiodic (1/f slope) activity and periodic activity (power in narrowband peaks) for each participant, as previously described^49^. Within the periodic data, we compared the total number of narrowband peaks, the central frequency of each peak, and power at that frequency across groups. Within the aperiodic data, we compared exponent and offset within each group. Our aims were three-fold: first, to examine the effect of stroke on the power spectrum across age; second, to explore how lesion location affects the power spectrum in the subset of participants with stroke; and finally, to examine the relationship between aperiodic activity and functional outcome in aged and young individuals after stroke.

## 2. Methods

### Data Sources

EEG data were obtained from previously collected data sets. Data from neurologically intact control data was from four different sources. Three were collected at the University of Utah with the involvement of two authors [KRL and MJE]^44,50,51^ and one was from a publicly available dataset from the Max Planck Institute^52^. Resting EEG data for people with stroke were taken from a previous study involving one of the authors [SC]^36^. For functional assessment of the stroke group, included patients were those with unilateral stroke, without contracture, significant subluxation or pain, severe neglect or apraxia, impaired level of consciousness, or a significant language deficit as previously described^36^.

### EEG Recordings

The specific details of each dataset’s recording parameters have been previously published as referenced above. Recording parameters for each dataset are summarized in Table 1. EEGs from all datasets were recorded in the resting state. Subjects were instructed to stare straight ahead with eyes open and while holding their body still. Original EEGs were recorded for 2-16 minutes depending on the study. Longer EEGs were manually truncated to 5 minutes prior to processing. Scalp EEGs were collected via 32, 62, 64, or 256 channel EEGs (electrodes and amplifiers outlined in Table 1) and were sampled at 1000 Hz^36,44^, 1024 Hz^51^, or 2500 Hz^52^.

**Table 1.**
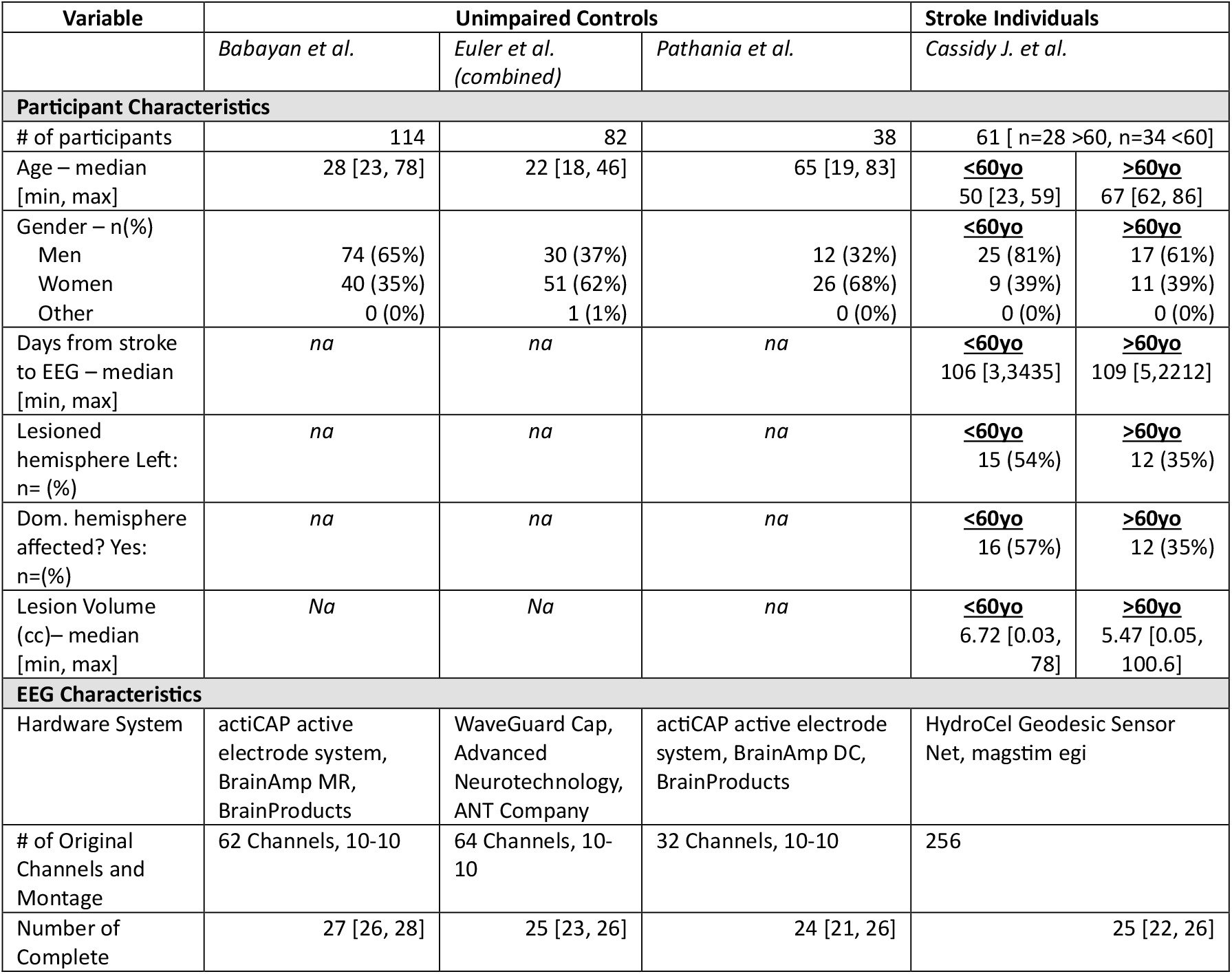
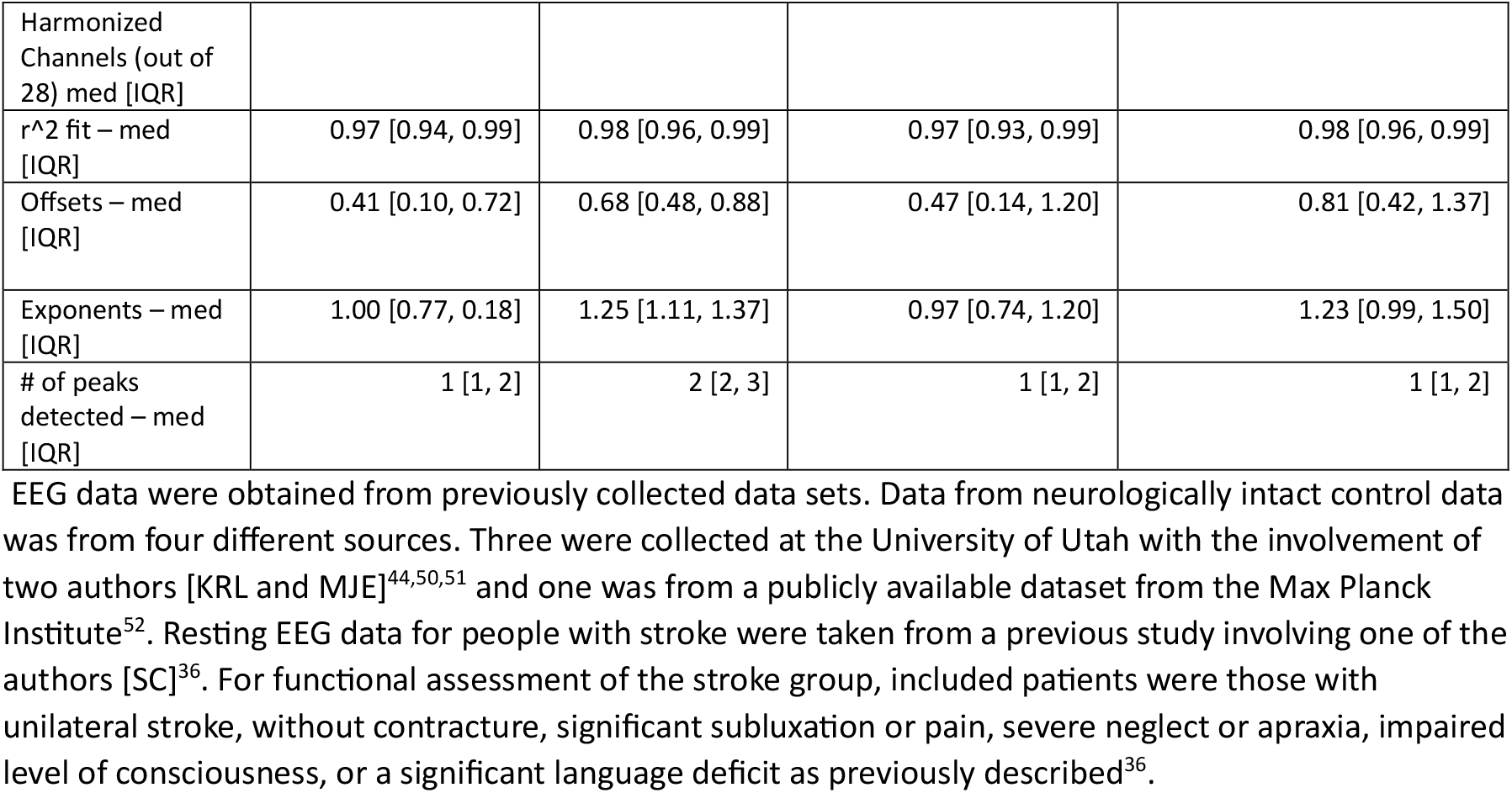
Demographic statistics and spectral parameterization fits for each sample.

| Variable | Unimpaired Controls |  |  | Stroke Individuals |  |
| --- | --- | --- | --- | --- | --- |
|  | *Babayan et al.* | *Euler et al.*<br>*(combined)* | *Pathania et al.* | *Cassidy J. et al.* |  |
| Participant Characteristics |  |  |  |  |  |
| # of participants | 114 | 82 | 38 | 61 [ n=28 >60, n=34 <60] |  |
| Age – median<br>[min, max] | 28 [23, 78] | 22 [18, 46] | 65 [19, 83] | <u><60yo</u><br>50 [23, 59] | <u>>60yo</u><br>67 [62, 86] |
| Gender – n(%) |  |  |  | <u><60yo</u> | <u>>60yo</u> |
| Men | 74 (65%) | 30 (37%) | 12 (32%) | 25 (81%) | 17 (61%) |
| Women | 40 (35%) | 51 (62%) | 26 (68%) | 9 (39%) | 11 (39%) |
| Other | 0 (0%) | 1 (1%) | 0 (0%) | 0 (0%) | 0 (0%) |
| Days from stroke<br>to EEG – median<br>[min, max] | *na* | *na* | *na* | <u><60yo</u><br>106 [3,3435] | <u>>60yo</u><br>109 [5,2212] |
| Lesioned<br>hemisphere Left:<br>n= (%) | *na* | *na* | *na* | <u><60yo</u><br>15 (54%) | <u>>60yo</u><br>12 (35%) |
| Dom. hemisphere<br>affected? Yes:<br>n=(%) | *na* | *na* | *na* | <u><60yo</u><br>16 (57%) | <u>>60yo</u><br>12 (35%) |
| Lesion Volume<br>(cc)– median<br>[min, max] | *Na* | *Na* | *na* | <u><60yo</u><br>6.72 [0.03,<br>78] | <u>>60yo</u><br>5.47 [0.05,<br>100.6] |
| EEG Characteristics |  |  |  |  |  |
| Hardware System | actiCAP active<br>electrode system,<br>BrainAmp MR,<br>BrainProducts | WaveGuard Cap,<br>Advanced<br>Neurotechnology,<br>ANT Company | actiCAP active electrode<br>system, BrainAmp DC,<br>BrainProducts | HydroCel Geodesic Sensor<br>Net, magstim egi |  |
| # of Original<br>Channels and<br>Montage | 62 Channels, 10-10 | 64 Channels, 10-10 | 32 Channels, 10-10 | 256 |  |
| Number of<br>Complete | 27 [26, 28] | 25 [23, 26] | 24 [21, 26] | 25 [22, 26] |  |

|  |  |  |  |  |
| --- | --- | --- | --- | --- |
| Harmonized Channels (out of 28) med [IQR] |  |  |  |  |
| $r^2$ fit – med [IQR] | 0.97 [0.94, 0.99] | 0.98 [0.96, 0.99] | 0.97 [0.93, 0.99] | 0.98 [0.96, 0.99] |
| Offsets – med [IQR] | 0.41 [0.10, 0.72] | 0.68 [0.48, 0.88] | 0.47 [0.14, 1.20] | 0.81 [0.42, 1.37] |
| Exponents – med [IQR] | 1.00 [0.77, 0.18] | 1.25 [1.11, 1.37] | 0.97 [0.74, 1.20] | 1.23 [0.99, 1.50] |
| # of peaks detected – med [IQR] | 1 [1, 2] | 2 [2, 3] | 1 [1, 2] | 1 [1, 2] |

### EEG Processing

Raw EEG data were imported into Matlab (Mathworks, Portola Valley, CA) and preprocessed as below using custom written Matlab code in conjunction with EEGlab (Swarz Center for Computational Neuroscience, UCSD, San Diego, CA). The data were down sampled to 250Hz and high pass filtered at 0.1Hz. Electrode data were re-referenced to the average reference using the ‘pop-reref’ function from EEGLab. The PREP pipeline^53^ was applied to reject excessively noisy or bad channels. The 50-60Hz line noise was removed with the Zapline-plus toolbox within EEG lab. Data were then decomposed into independent components using the ‘runica’ function to classify signals as cardiac rhythms, muscle artifacts, ocular movement or blinks, and then subsequently removed from the EEG signal using blind source separation via the AAR plugin for EEGlab (https://github.com/germangh/eeglab_plugin_aar/blob/master/README.md). As a final pre-processing step, any flatline channels, low-frequency drifts, noisy channels, short-time bursts and incompletely repaired segments from the data were removed with the clean_rawdata plugin for EEGlab (https://github.com/sccn/clean_rawdata). After undergoing artifact rejection, processed EEG data from all datasets were then subjected to fast Fourier transform using a Hamming window with 50% overlap between segments. The power spectral density was estimated using Welch’s method with the number of frequency bins varying according to the sampling rate. This was performed at each channel for each participant. All code is available here: https://github.com/margareteisfelder/Automated-EEG-Cleaning-Pipeline/blob/main/README.md.

### Spectral Parameterization

Following initial processing, spectral analysis was performed using the *specparam* toolbox^40^ running in Python (version 3). At each electrode, spectra were decomposed into periodic and aperiodic components (2-25 Hz). Settings for the specparam algorithm were as follows: peak width limits of 0.5-12, 6 maximum peaks, a minimum peak height of 0.1, and a peak threshold of 2. The quality of the of the parameterization was assessed via r^2^ values provided by the algorithm and summarized for each data set in Table 1.

As shown in Figure 1, at each electrode we extracted two *aperiodic*, broadband components – the offset and the exponent – and a number of *periodic*, narrowband components for every Gaussian peak that was identified – the central frequency of the Gaussian, the peak power at the central frequency, and the full width half maximum of the Gaussian at that central frequency.

**Figure 1.**
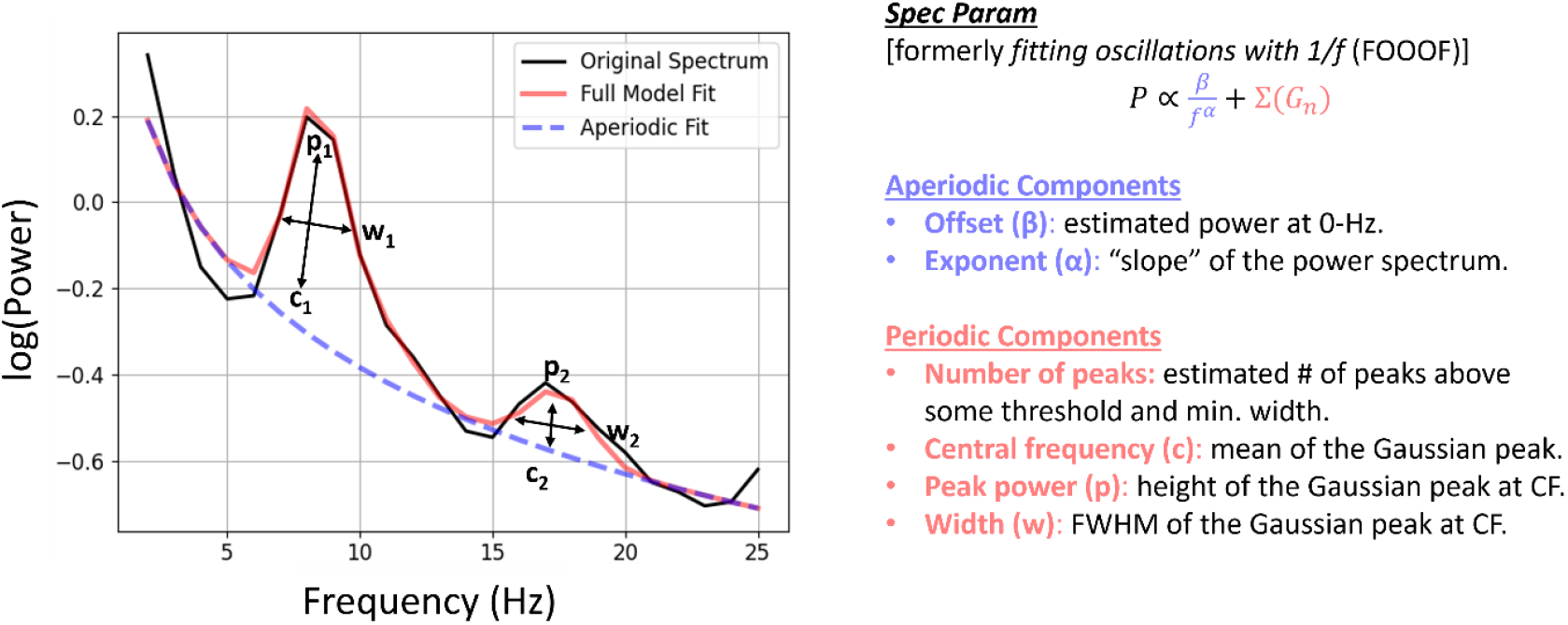
Illustration of the spectral parameterization (spec param) algorithm fit to data from a sample electrode from a control subject. At every electrode for every subject, we extract two aperiodic components (the offset and the exponent) and a number of Gaussian peaks (Gn). For every Gaussian, we obtain the central frequency, the peak power at that central frequency, and the full-width half-maximum (FWHM) of the Gaussian at the central frequency.

We also faced a challenge harmonizing the EEG data across these different samples owing to the different number of channels and montages used in each original data collection, see Table 1. However, we did have 28 channels common to all datasets that could be harmonized and included in statistical analyses. Note that for the high density EEG from the stroke dataset- we had to use the HydroCel geodesic sensor nets approximate 10 – 10 equivalents (see: https://www.egi.com/images/HydroCelGSN_10-10.pdf), but to the best of our ability, these 28 channels come from the same location on the scalp in all samples: “F3”, “Fz”, “F4”, “F7”, “F8”, “FC1”, “FC2”, “FC5”, “FC6”, “C3”, “Cz”, “C4”, “CP1”, “CP2”, “CP5”, “CP6”, “P3”, “Pz”, “P4”, “P7”, “P8”, “O1”, “Oz”, “O2”.

The r^2^ value, (representing how well the model fits the EEG power spectral data) across all channels was generally quite high: median=0.97, IQR=[0.94, 0.99] in healthy controls, median=0.97, IQR=[0.96, 0.99] in adults with stroke. Further, the r^2^ value was generally quite high regardless of scalp location, suggesting that our processing pipeline was suitable for capturing variation in the EEG power spectrum across regions and populations: frontal median=0.98, IQR=[0.95, 0.99], central median=0.94, IQR=[0.94, 0.99], parietal median=0.97, IQR=[0.94, 0.99], and occipital median=0.98, IQR=[0.95, 0.99]. We created grand average spectra for both the control groups and the stroke groups (Figure 3). Similar to our previous work, these figures combined with our r^2^ values suggest that appropriate parameters were chosen, adequately capturing both periodic and aperiodic variation across the scalp and between participants.

### Magnetic Resonance Imaging

Magnetic Resonance Imaging (MRI) data was available from the stroke data set and have been previously described^36^ In brief, high resolution T1-weighted images were acquired with on a Philips Achieva 3T MRI scanner using 3D magnetization-prepared rapid gradient echo (MPRAGE) Parameters are as follows: Repetition time=8.5ms, echo time=3.9 ms, slices=150. Voxel size- 1 × 1 × 1mm^3^. Infarct volumes were previously outlined by hand on all T1 weighted MRI images. Individual subject MPRAGE images were linearly aligned to a reference atlas (MNI152) using *flirt* in FSL. The transform matrix for the subject-to-atlas alignment was then used to transform the stroke mask into atlas space. Stroke masks in atlas space were combined across subjects to produce a stroke frequency map representing the probability that each voxel was within the stroke mask for any individual subject (Figure 2).

**Figure 2.**
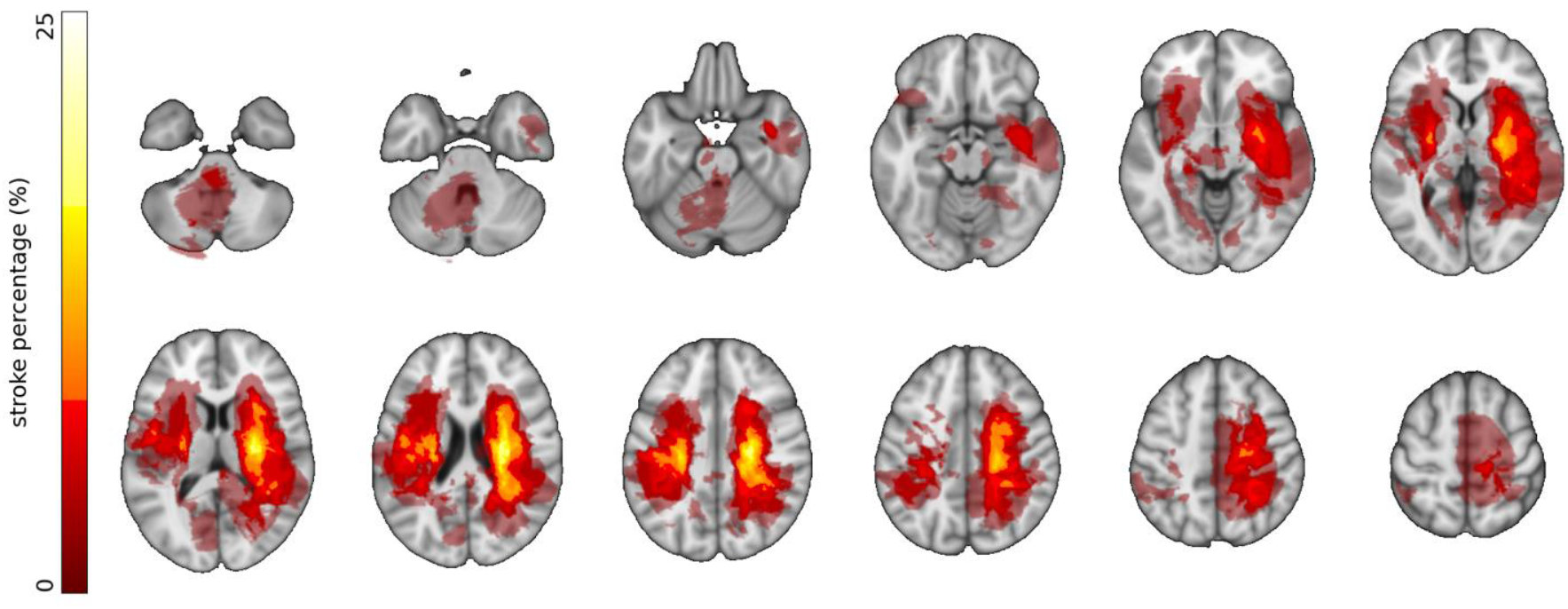
Infarct Masks from 61 patients overlaid on T1-weighted images. Brighter colors correspond to increased frequency of damage in a voxel. Right side of image corresponds to anatomical right so as to align with EEG presentations.

### Functional Assessment

Previously obtained functional assessment scores (Box and Block Test) were utilized for our analysis. Upper extremity motor deficits were assessed via one-time scoring on the Box and Block test measured on the same day as the EEG recording, during which subjects were asked to transfer as many blocks as possible from one side of a box to another in 60 seconds using the affected arm. Higher scores (number of blocks transferred) correspond to better function in the paretic limb, as previously described by Cramer et al.^54-56^

### Statistical Analysis

Our statistical analysis can be delineated into two main sections: comparison of healthy controls to adults with stroke, and associations of the EEG components with clinical measures in the stroke subgroup. In each analysis, we ran a series of mixed-effect regression models^57,58^ to address hypotheses about the aperiodic offset of the EEG power spectrum, the number of Gaussian peaks detected, the central frequency of those peaks, and the power at those frequencies.

As we did not have strong hypotheses about the spatial location of the signal, and there is not a clear way to map three dimensional locations/volumes onto EEG scalp topography, we treated subject, specific electrode, and region (defined as F, C, P, or O electrodes) as random effects. This approach allows us to account for multiple sources of variation that contribute to the periodic and periodic signals, reducing statistical dependence in the errors of the model^59,60^. Conceptually, this means that models can be thought of as scalp average associations, with regions and electrodes treated as a random sample of all possible locations in the same way subjects are a random sample of the population.

In the stroke subsample, models include a fixed effect of electrode hemisphere (contralesional or ipsilesional) as all participants had unilateral stroke. Central electrodes (Fz, Cz, Pz, and Oz) were excluded from these analyses so that all electrodes could be defined as either ispilesional or contralesional. All models also controlled for age, sex, lesion volume, and time from stroke to the EEG data collection.

When plotting the relationship between the box and block test (BBT; a clinical measure of upper extremity function) and the aperiodic slope, we noticed a potentially nonlinear affect of age on the relationship. Therefore, in that model, rather than treating age continuously, we included age group as a categorical factor (<60 versus ≥60) and included interactions of age group with BBT, region, and hemisphere when predicting the aperiodic exponent.

Approximate normality of the residuals and random effects was confirmed with visual inspection of quantile-quantile normal plots, but note that mixed-effect models are also generally robust to violations of normality^61^. Outcomes for the number of peaks and the power for each peak were square root and log-transformed, respectively, prior to analysis to achieve approximate normality of residuals.

Coefficients and 95% confidence intervals from these analyses were then back transformed into their original units. P-values were calculated using the Welch-Satterthwaite approximation for the degrees of freedom^62^ and the statistical significance for all analyses was defined as α=0.05. Given the early phase nature of this work, we considered the cost of Type II errors (“misses”) to be high relative to Type I errors (“false alarms”) and did not apply a correction for multiple comparisons^63^. All analyses are reported (see supplemental appendices for all models) with p-values reported continuously, so that readers can make more conservative/corrected judgments if desired, and p-values are complemented by estimates of effect size and uncertainty^64^.

We performed a data-driven analysis to describe the multivariate spatial relationship between stroke location and periodic scalp-measured electrical activity. Canonical correlation is the multivariate generalization of the more familiar pairwise correlation (e.g., Pearson)^65^. Canonical correlation identifies linear combinations (topographies) of one variable (stroke location) that is maximally correlated with the topography of another variable (exponent (α) at a given electrode). To reduce variability, increase interpretability, and given the lack of an *a priori* laterality hypothesis, we transformed the images such that all strokes were on the same side. This was performed by identifying strokes with center of mass on the contralateral side and reflecting the lesion mask across the mid-sagittal plane and the α values across the mid-sagittal electrodes. The stroke mask matrix (dimensions = subjects × voxels) and α matrix (dimensions = subjects × electrodes) were subjected to canonical correlation analysis. Given the high dimensionality of the data, we used a sparse algorithm well suited for poorly conditioned data^66^. The first canonical correlation component (i.e., corresponding to the largest eigenvalue) comprises a spatial stroke mask and EEG weights. The original data, when projected onto these weights, result in maximal correlation. In this way, canonical correlation is a data driven tool for describing multivariate relationships within these data.

## RESULTS

We divided our analysis into 3 sections. First, we broadly examined the influence of stroke and age on the power spectrum as a whole. Next, we examined the influence of lesion location on the aperiodic and periodic components of the power spectrum. Finally, we examined the relationship between aperiodic activity and motor outcomes (box and block testing) within the stroke group.

### Age- and Stroke-Related Changes in the Periodic and Aperiodic Components of the Power Spectrum

Prior work has demonstrated broad changes in power spectral density after stroke^33^. We first examined whether our data also demonstrated similar changes between the stroke and non-stroke groups. Given other work showing significant spectral changes with aging^31^, as well as the unique relevance of stroke to the aged brain^5^, we further examined spectral density across aged and young groups with and without stroke. We generated average power spectral densities for each patient (average of all electrodes across the scalp) and averaged across groups (Young patients <60 years, Aged Patients >60 years, and stroke patients within each group). Spectral density comparisons for each group are illustrated in Figure 3.

**Figure 3.**
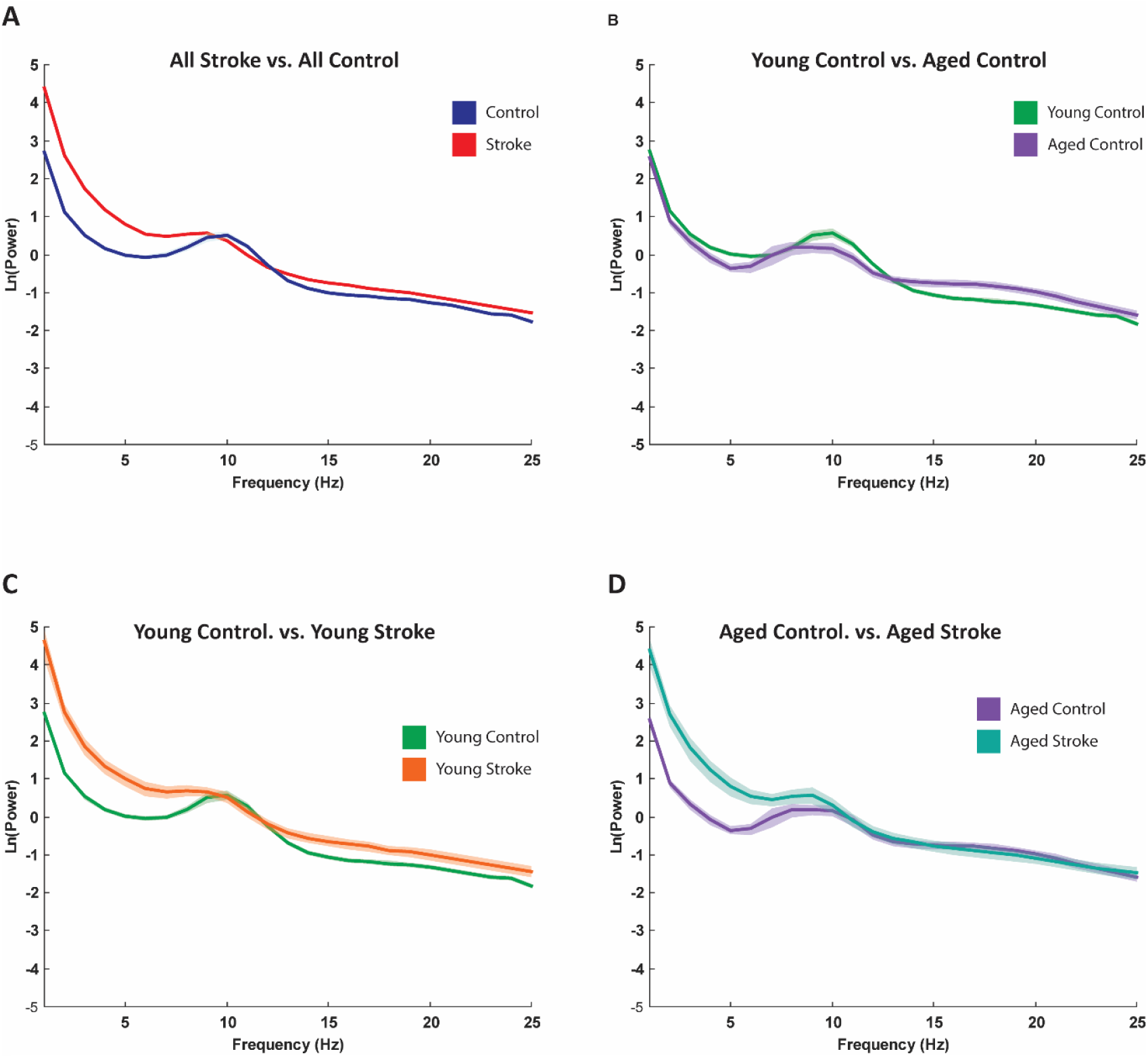
Power spectral density as generated by the specparam algorithm averaged across the scalp and across groups. Comparison of all control to all stroke individuals. B. Comparison of Aged vs. Young Groups in non-stroke individuals. C. Comparison of Young control vs. Young Stroke invididuals. D. Comparison of Aged Control vs. aged stroke individuals.

Consistent with work from other groups^39,44^, we observed decreased low frequency power in the aged group (Figure 3b) when compared to the young group. Also consistent with prior studies^35,67,68^, stroke patients were noted to have higher power at slower frequencies (Figure 3a). After broad spectral visualization we focused on individual differences within the periodic and aperiodic components of the power spectrum.

#### 1. Age and Stroke-Related Changes in the Aperiodic Component of the Power Spectrum

Given recent data showing that stroke-related changes in spontaneous neuronal activity are partially driven by changes in aperiodic activity^45^, as well as work showing age related changes in aperiodic activity and the relationship between this activity and cerebral function^44,49^, we examined age related changes in spectral slope (quantified as exponent) in both the stroke and non-stroke groups.

##### 1.a Exponent

As seen in Figure 4, there were main-effects of Age, F(1,295)=28.5, p<0.001, and Group (stroke vs. neurotypical), F(1,295)=67.3, p<0.001, and a statistically significant Group x Age interaction, F(1, 295)=4.45, p=0.036. In healthy controls, we saw a characteristic flattening of the power spectrum with increased age, β=-9E-3, 95%CI = [−10E-2, −7E-3]. This flattening has been previously observed by our group^44^ and others^39^ and it has been hypothesized that these changes may reflect age related changes in the balance of excitatory vs. inhibitory neuronal activity^39^. In adults with stroke, however, the observed pattern of progressive age-related flattening of slope was significantly attenuated, β=-4E-3, 95% CI = [−8E-3, −6E-4]. For the full mixed effect model, see Supplemental Table i. As seen in Figure 4, there are some data to suggest that this slope differs across the scalp. In our model, however, we treated region as a random-effect to get a scalp-wide estimate of differences in the exponent due to age (i.e., treated region as a random sample of possible regions). We explore regional differences in the exponent in the stroke sub-sample below.

**Figure 4.**
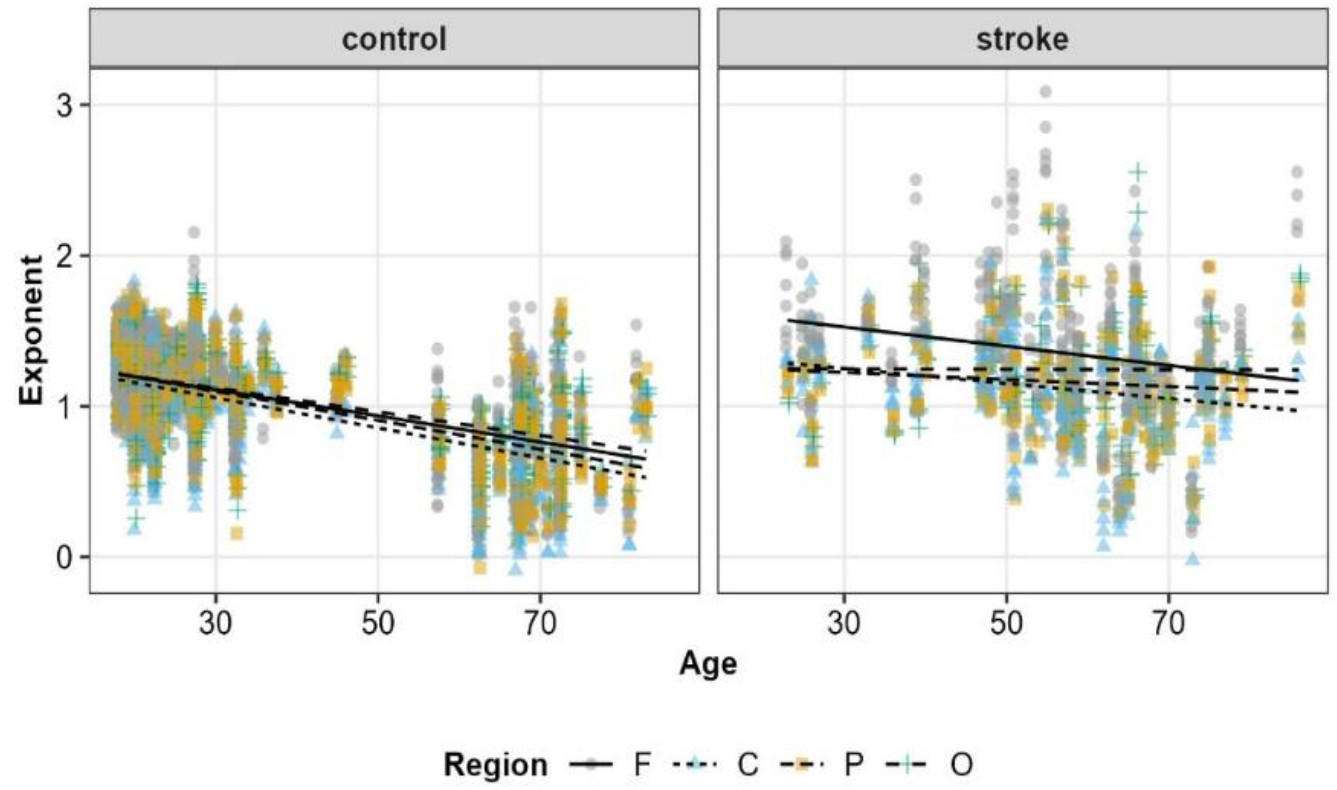
Aperiodic exponents from the spectral parameterization algorithm as a function of age and scalp region (F=Frontal, C=Central, P=Parietal, O=Occipital) for neurotypical controls (left) and adults with stroke (right). Lines show the OLS regression slope for exponent ∼ age in that region

To better understand this Group x Age interaction, we divided age into categories based on the distribution of the stroke sub-sample. For adults 18-36 years old, the mean exponent for healthy controls was 1.15, 95% CI=[1.10, 1.21], and for adults in this age group with stroke was 1.29, [1.10, 1.48], p_difference_=0.177. For adults 37-59, the mean exponent for controls was 0.94, [0.75. 1.13], and for adults with stroke was 1.30, [1.19, 1.40], p_difference_< 0.001. For adults 60-74, the mean exponent for controls was 0.73, [0.64, 0.81], and for adults with stroke was 1.08, [0.97, 1.19], p_difference_<0.001. And for adults 75-86, the mean exponent for controls was 0.77, [0.56, 0.97], and for adults with stroke was 1.34, [1.13, 1.54], p_difference_<0.001.

#### 2. Age and Stroke-Related Changes in the Periodic Component of the Power Spectrum

Longstanding evidence has demonstrated disruptions in periodic activity after stroke^33^. These disruptions have been correlated to outcomes after stroke^69^. We therefore also examined age and stroke-related changes in the periodic component of the spectrum. To better quantify changes within the periodic component of our data, we first observed the number of distinct (Gaussian) peaks isolated by the model within each group, then further compared two aspects of identified periodic activity within groups: the central frequency at each of those peaks and the power at that frequency within each group.

##### 2.a Number of Peaks

A mixed-effect model examining the number of peaks identified by the spectral parameterization algorithm, Supplemental Table ii, showed main-effects of Age, F(1,291)=6.42, p=0.012, and Group, F(1,291)=20.27, p<0.001, and a Group x Age interaction, F(1,291)=5.67, p=0.018. Breaking this down by age groups, for adults 18-36, healthy controls had a mean number of peaks = 3.1, 95% CI=[2.9, 3.3], and adults with stroke had a mean = 2.0, [1.6, 2.6]. For adults 37-59, controls had a mean of 2.7 peaks, [2.2, 3.3] and adults with stroke mean = 2.3, [1.9, 2.5]. For adults 60-74 years, controls had a mean = 2.4, [2.2, 2.6], and adults with stroke mean = 2.1, [1.8, 2.4]. For adults 75-86 years, controls had a mean = 2.5, [2.0, 3.2], and adults with stroke mean = 2.0, [1.5, 2.6]. Thus, the algorithm generally identified more peaks in the controls compared to adults with stroke, but this difference tended to be larger in younger adults.

##### 2.b Central Frequency

Age associated changes in center frequency of periodic activity have been previously reported^70,71^. We therefore examined the relationship between central frequency of periodic activity across identified peaks within age groups in the stroke and healthy control groups (Figure 5A). When looking at the central frequency of the identified peaks, as shown in Figure 5, there was a (tautological) effect of Band, F(2, 13343)>100, p<0.001, but also main effects of Age, F(1, 454)=7.66, p=0.006, and Group, F(1, 439)=4.37, p=0.037, with significant interactions of Band x Age, F(2, 13353)=52.0, p<0.001, and Group x Band, F(2, 13342)=3.07, p=0.046. None of the other effects were statistically significant (p>0.327). The full mixed-effects model is shown in Supplemental Table iii. To better understand the Age x Band interaction, we fit a model with Age, Group, and their interaction in each canonical frequency band, except for the delta band because few peaks were identified in that range.

**Figure 5.**
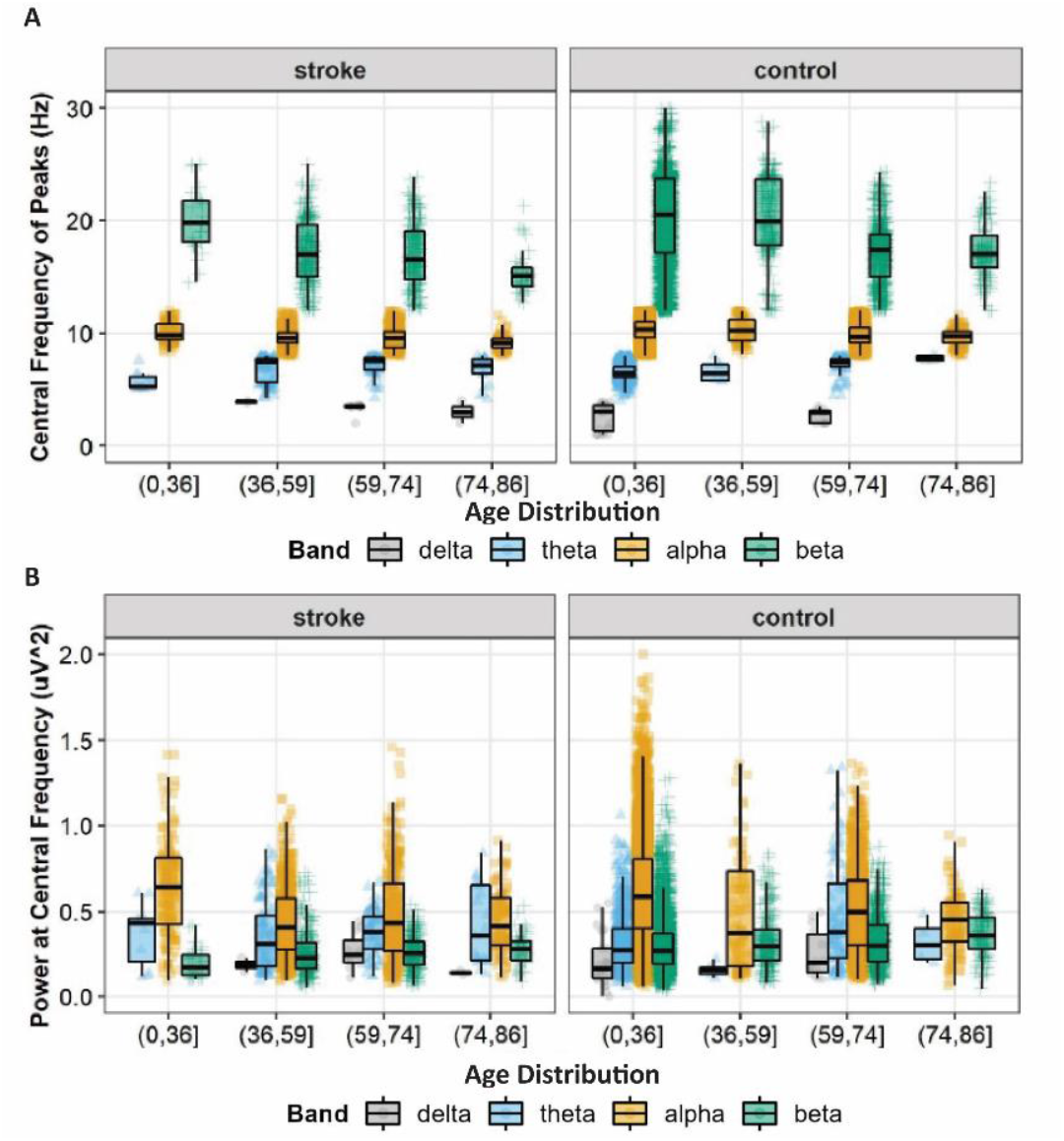
Central frequency of identified peaks (A) and the power at the central frequency (B) as a function of age and the canonical frequency bands of the EEG power spectrum.

In the theta band, there was a statistically significant effect of Age, F(1,171)=5.14, p=0.025, but no significant effect of Group, F(1,174)=2.25, p=0.135, nor was there an Age x Group interaction, F(1,171)=0.18, p=0.676.

In the alpha band, there were main-effects of Age, F(1,280)=10.00, p=0.002, and Group, F(1, 281)=6.66, p=0.010, but not an Age x Group interaction, F(1, 280)<0.01, p=0.882. On average, younger adults had higher central frequencies for peaks identified in the alpha range, β=-1E2, 95% CI=[−02E2, −4E3]. Adults with stroke also tended to have lower central frequencies for alpha peaks, mean=9.70, [9.49, 9.91], compared to healthy controls, mean=10.00, [9.87, 10.14].

In the beta band, there as a main-effect of Age, F(1,352)=18.77, p<0.001, but not a main-effect of Group, F(1,328)=2.10, p=0.148, nor an Age x Group interaction, F(1,353)=0.93, p=0.334. On average across groups, younger adults had higher central frequencies for peaks identified in the beta range, β=-0.05, 95% CI=[−0.08, −0.03].

##### 2.c Power at Central Frequency

Changes in the power of defined frequency bands with both aging and stroke have been previously identified and correlated to stroke outcomes and cognitive function^31,33^. We therefore examined the relationship between power at our identified central frequencies within the aged healthy control and stroke groups. Power at the central frequency, shown in Figure 5, varied as a function of Band, F(2, 13313)>100, p<0.001, and Group, F(1, 313)=4.33, p=0.038. There was not a statistically significant main effect of Age, F(1, 318)=0.60, p=0.439, but there were significant interactions of Band x Age, F(2, 13306)=45.96, p<0.001, and Group x Band, F(2, 13306)=9.5, p<0.001. None of the other effects were statistically significant (p>0.188). The full mixed-effects model is shown in Supplemental Table iv. As with central frequency, we followed these interactions with smaller models focused on each band, but again the delta band was excluded from analyses due to the small number of observations in that range.

In the theta band, there were no statistically significant effects of Age, F(1, 175)=1.89, p=0.177, Group, F(1,178)=0.05, p=0.816, nor a Group x Age interaction, F(1,176)<0.01, p=0.980.

In the alpha band, there was a statistically significant effect of Age, F(1, 283)=7.13, p=0.008, but not a statistically significant effect of Group, F(1, 284)=3.01, p=0.083, nor an Age x Group interaction, F(1, 283)=0.13, p=0.715. On average, alpha power was reduced in older participants compared to younger participants, β=-2E3, 95% CI=[−3E3, −5E4].

In the beta band, there was a statistically significant effect of Age, F(1, 349)=5.01, p=0.026, and Group, F(1, 326)=24.46, p<0.001, but not an Age x Group interaction, F(1, 349)=2.00, p=0.158. On average, beta power was increased in older participants compared to younger participants, β=9E4, 95% CI=[1E4, 2E3].

Healthy controls also tended to have higher beta power, mean=1.29, 95% CI=[1.26, 1.32], then adults with stroke, mean=1.19, [1.14, 1.24].

### Spatial Relationships within the Stroke Subgroup

#### 1. Influence of Lesion Location and Size on the Aperiodic Component of the Power Spectrum

Stroke location^72,73^ and lesion volume^74^ each have a significant impact on cerebral function. Additionally, significant work has described the presence of diaschisis or widespread influence of stroke beyond the infarcted region in stroke pathology and recovery^75^. We therefore examined the relationship that spectral content has with lesion location (as defined by broad brain region) and lesion volume within the stroke group.

Recent work has shown that stroke is associated with a significantly greater exponent in both the ipsilesional and contralesional hemisphere^45^. We sought to add to this work by examining for differences in aperiodic exponent within electrodes in broad cortical regions (frontal, central, parietal, occipital) of the ispsilesional or contralesional hemispheres. We then sought to more precisely map the relationship between aperiodic exponent and lesion location by using MRI and EEG topgography. We also examined for association between lesion volume and exponent overall. We did not observe a significant relationship in our data between lesion seize and aperiodic exponent.

##### 1a. Influence of Lesion Hemisphere on Aperiodic Exponent Within Broad Cortical Regions

Controlling for participants’ gender, age, and days from stroke to EEG recording, there were significant main effects of hemisphere (contralesional or ipsilesional), F(1,70)=7.79, p=0.007, and channel Region, F(3,15)=14.82, p<0.001, and a Hempsipher x Region interaction, F(3, 912) = 7.90, p<0.001. (Full details in Supplemental Table v.) In the frontal region, contralesional electrodes had a mean exponent = 1.32, 95% CI=[1.21, 1.43], and ipsilesional electrodes had a mean = 1.29, [1.18, 1.4], p_diff_=0.290. In the central region, contralesional electrodes had a mean exponent = 1.06, [0.95, 1.17], and ipsilesional electrodes had a mean = 1.12, [1.00, 1.23], p_diff_=0.023. In the parietal region, contralesional electrodes had a mean exponent = 1.08, [0.96, 1.20], and ipsilesional electrodes had a mean = 1.18, [1.06, 1.30], p_diff_=0.002. And finally in the occipital region, contralesional electrodes had a mean exponent = 1.18, [1.04, 1.31], and ipsilesional electrodes had a mean = 1.26, [1.12, 1.40], p_diff_=0.022. Thus, ipsilesional electrodes generally tended to have steeper slopes than contralateral electrodes, with the exception of a non-significant difference in the frontal electrodes.

##### 1b. Influence of stroke location from MRI on Aperiodic Exponent

As noted in Figure 6, we observed a significant relationship between stroke hemisphere and aperiodic activity (exponent). This analysis is limited to hemispheric regions. We next sought to more precisely quantify the relationship between exponent and lesion location using MRI data from the participants within the stroke group.

**Figure 6.**
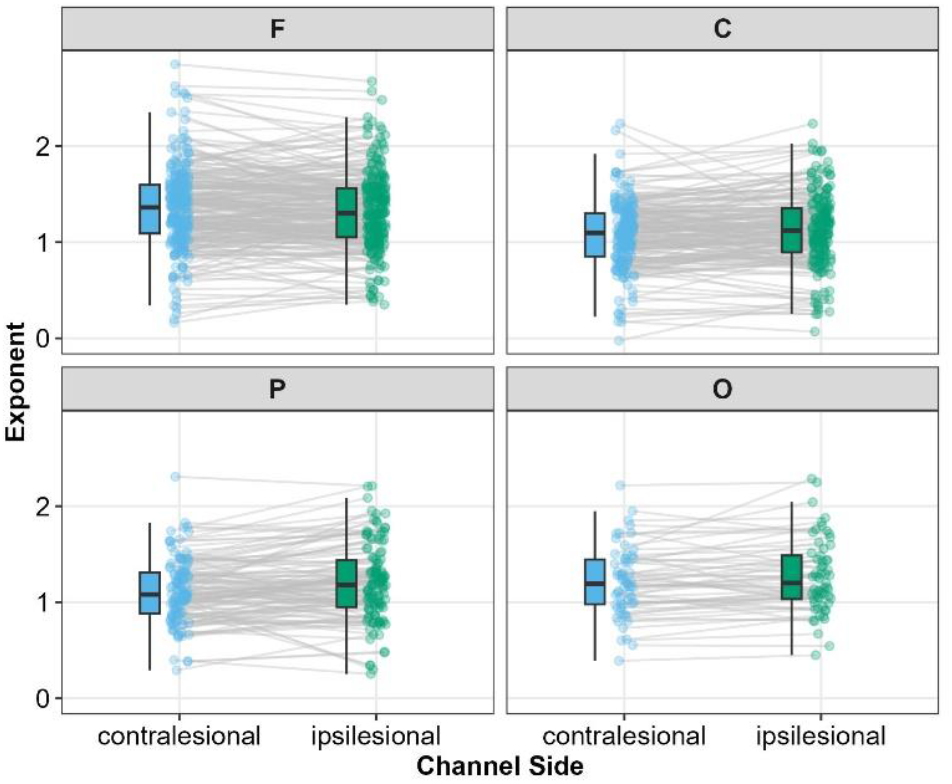
Aperiodic exponents as a function of channel region (F: Frontal, C: Central, P: Parietal, or O: Occipital) and channel side (either ipsilesional or contralesional).

We used canonical correlation to examine the multivariate relationship between stroke topography and topography of disrupted EEG as measured by the exponent (α) (Figure 7). The first canonical correlation is defined by a stroke topography that is maximally correlated with an EEG topography. Recall that, absent an *a priori* laterality hypothesis, we transformed images into stroke side vs. non-stroke side instead of anatomical left vs. right. The first canonical correlation identified a stroke topography maximally loaded in the deep white matter, which also encompasses deep grey nuclei. In contrast, stroke topography was negatively loaded in the contralesional brain. The EEG topography was maximally loaded ispilesionally with maximal values in frontal and pre-frontal electrodes. This demonstrates that the presence of a stroke in the deep structures of the brain is positively related to changes in EEG α over the ipsilateral frontal scalp with minimal impact on the contralateral EEG activity. This indicates that deep strokes manifest as steeper EEG spectra over the affected side.

**Figure 7.**
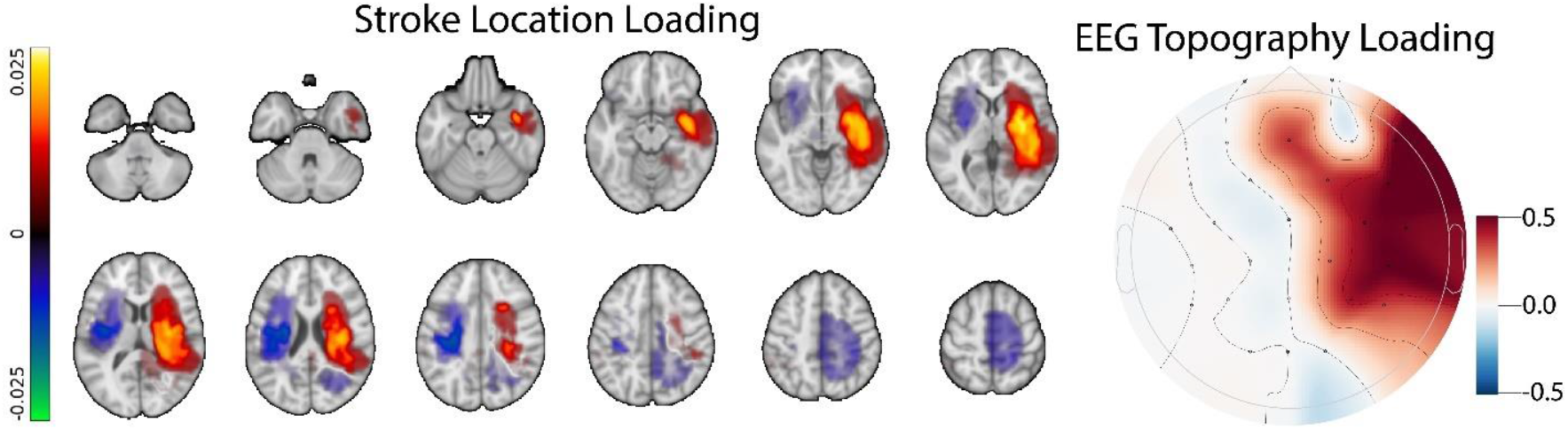
Stroke spatial topography (left) and EEG α topography (right) corresponding to the first canonical correlation between stroke location and EEG \beta coefficient. N.B. that strokes were aligned so that they all appear on the right; the EEG was similarly transposed across the mid-sagittal plane to define a stroke side (right) and non-stroke side (left). This result shows a strong correlation between deep strokes and increased slope of the EEG signal over ispilesional frontal electrodes.

#### 2. Influence of Lesion Location on the Periodic Component of the Power Spectrum

Stroke is associated with region specific changes in periodic activity-particularly within the contralateral cortex^76^. We therefore examined the influence of hemisphere (ipsilesional vs. contralesional) on the periodic components of the spectrum.

##### 2a. Number of Peaks and Central Frequency

There were no statistically significant differences in the number of peaks or the central frequencies of peaks as a function of ipsilesional vs. contralesional hemisphere, see Supplemental Tables vi and vii. Thus, the spectral parameterization algorithm appeared to be identifying peaks on the contralesional and ipsilesional side in similar ways, and we can more robustly interpret differences in power for the observed peaks.

##### 2.a Narrowband Power at Central Frequency

Controlling for participants gender, age, and days from stroke to EEG recording, there were statistically significant main effects of Band, F(2,1377)=218.81, p<0.001, and Region, F(3,28)=18.10, p<0.001, on narrow-band power. There were also statistically significant interactions of Band x Channel Region F(6,1345)=2.96, p=0.007, and Hemisphere x Region, F(3, 1364)=3.48, p=0.015. No other effects were statistically significant (p’s>0.070, see Supplemental Table viii. Post-hoc tests of these interactions revealed that narrowband power was generally lower at the frontal electrodes compared to central and parietal electrodes, and these effects were more pronounced on the ipsilesional side: i.e., contralesional F – C, p=0.013; F – P; p=0.002; F – O, p=0.888. Ipsilesional F – C, p<0.001; F – P, p<0.001; F – O = 0.017. Further, power was generally highest in the alpha band, but the strength of this effect depended on the specific region of the scalp. As with previous analyses, the delta band was excluded due to a limited number of peaks identified in the delta band.

### Relationship Between Aperiodic Activity and Motor Outcomes

Changes in canonical periodic activity have been extensively examined as potential prognostic biomarkers after stroke (Reviewed by Sutcliffe L et al)^77^. We sought to examine whether abnormalities in aperiodic activity (recently shown to contribute to post-stroke spectral slowing^45^) are associated with motor outcomes after stroke. We examined the relationship between exponent and Box and Block scores in the stroke group. We observed a relatively linear correlation between performance on Box and Block testing and exponent, with steeper slopes being associated with better performance (Figure 9).

**Figure 8.**
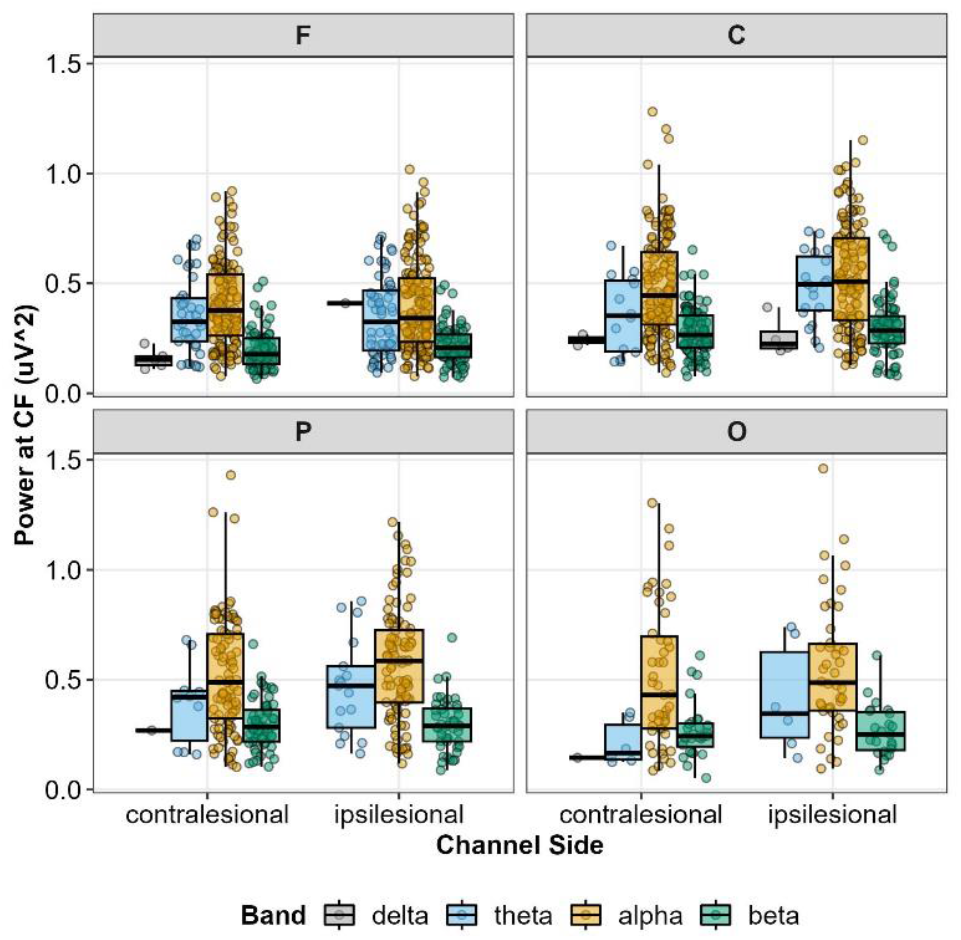
Narrowband power as a function of the canonical frequency bands (delta = grey, theta = blue, alpha = gold, and beta = green), scalp region (Frontal, Central, Parietal, or Occipital), and channel side relative to the lesion (contralesional or ipsilesional).

**Figure 9:**
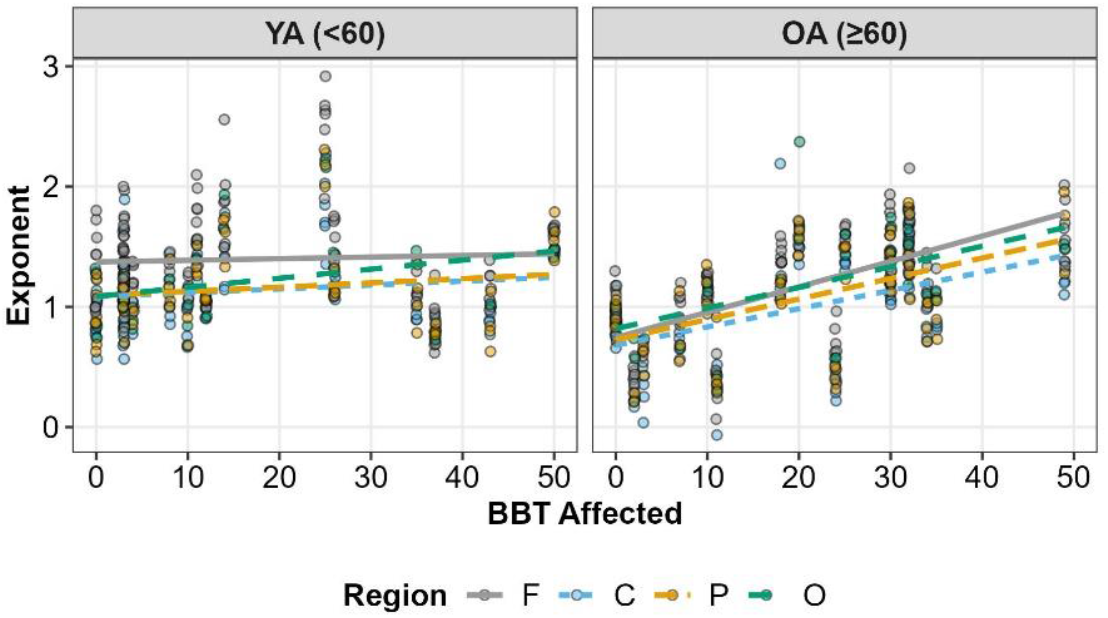
Aperiodic exponents as a function of age group, region, and box and block test (BBT) score (higher values = better performance). Note that data points show repeated observations within a subject (accounted for by the random effects of the model). Lines show simple ordinary least squares

We noted that young individuals with stroke showed similar exponents to those without stroke, but aged individuals with stroke demonstrated an overall higher exponent than those without (Figure 4). We therefore examined the relationship between exponent and performance on Box and Block testing in aged (>60yo) and young (<60) EEGs within the stroke group. Our mixed effect model associating aperiodic exponents with BBT scores showed a statistically significant main-effect of BBT, F(1,37)=8.50, p=0.006, and an Age Group x Region x BBT interaction, F(3, 566) = 3.27, p=0.021. As shown in Figure 9, the relationship between exponents and BBT scores was generally weaker for younger adults, with stronger relationships between exponents and BBT scores for older adults. Overall, greater exponent was associated with better performance, with this relationship being substantially more notable in older adults. However, the degree of this moderation also depended on the scalp region; being strongest over the frontal electrodes (Age Group x BBT p=0.045), weaker in the central (Age Group x BBT p=0.087) and parietal electrodes (Age Group x BBT p=0.082), and weakest over the occipital electrodes (Age Group x BBT p=0.439). See Supplemental Table ix for the full details of the model.

## DISCUSSION

Here we examined the differences in periodic and aperiodic activity in healthy young and old individuals as well as younger and older individuals with stroke. We first examined the influence of stroke and age on the periodic and aperiodic components of the power spectrum. In keeping with prior work, we observed a general decrease in low frequency power with age as well as increased power spectral density (particularly in low frequencies) after stroke (Figure 3). We then compared aperiodic activity (quantified as exponent of the aperiodic spectra) in all groups and noted a significant age-related decrease in slope (consistent with prior work)^45^ (Figure 4). We further noted that the age-dependent decrease in spectral slope was significantly attenuated in individuals with stroke (Figure 4). Aged individuals with stroke had a significantly steeper spectral slope than aged control individuals. This disparity in slope was not observed in young individuals with stroke. Within the periodic component of the spectrum, we found that stroke was associated with fewer frequency-specific peaks, and this effect was greatest in young individuals. We also noted effects of both age and stroke on central frequency and power of periodic activity (Figure 5). We next examined the influence of lesion location and size on the periodic and aperiodic components of the power spectrum within the subset of subjects with stroke. We noted power spectra were steeper (i.e., larger exponents) on the ipsilesional hemisphere (Figure 6).

Interestingly, we also noted that deep lesions had a disproportionate impact on exponent (Figure 7). Finally, we examined the relationship between aperiodic slope and motor performance. We noted an inverse correlation between performance on box and block testing after stroke and slope with higher exponent values (steeper slope) being associated with better performance (Figure 9a). After subdividing the stroke group into young and old individuals, we found this relationship to be dependent on age.

Within younger stroke patients, we noted no correlation between exponent and functional performance however, within the older group, there was a clear relationship between exponent and box and block performance. Older patients with greater slope performed better on functional assessment (Figure 9b).

Together, these results indicate spatial and age dependent relationships between stroke aperiodic cortical activity. Furthermore, these results suggest that age specific changes in periodic activity are correlated to stroke motor outcomes (box and block testing).

Prior work has demonstrated several EEG changes associated with stroke. Stroke is classically associated with spectral slowing characterized by diminished high frequency (beta/gamma) activity^78^ as well as increased low frequency power^35,67,68^. Delta (1-4HZ) power in particular has been reported to correlate with both injury size and motor recovery^36^. Recent work demonstrated that “spectral slowing” observed after chronic stroke is not merely a reflection of changes in periodic activity, but rather a combination of periodic shifts and changes in aperiodic activity (spectral slope)^45^. This work suggested that increased low frequency power observed after stroke was at least partially explained by a shift in aperiodic activity toward lower frequencies as quantified by greater aperiodic exponent. This work built on prior animal work suggesting that increased low power activity was a reflection of a change in aperiodic activity and correlated to outcomes in a rat model of stroke^79^. Our data support these findings with a similar observation that stroke is associated with significant changes in aperiodic activity as reflected by a greater aperiodic exponent in individuals with stroke. Here we note a significant age-dependent effect on aperiodic activity with young individuals after stroke having similar exponents to those without stroke while older adults with stroke have much higher exponents than those without stroke. This change in exponent is especially striking as it is the opposite of the typical direction of exponent change with aging. Together, these findings raise the intriguing possibility that stroke-associated changes in aperiodic activity may reflect a unique pathology of aging. It is well-established that older individuals with stroke have worse outcomes^80^ and diminished functional recovery^81^. It is interesting to speculate that age-specific disruptions in neuronal activity may contribute to diminished plasticity and recovery in aged individuals after stroke. This is particularly intriguing given our finding that higher slopes correlate with better functional performance in older individuals but not younger individuals.

As others have noted, the physiologic underpinnings of aperiodic activity and aperiodic slope in spectral parameterization of EEG recordings are still being elucidated, however, a significant body of evidence has suggested that these slope changes may reflect changes in the balance of network level excitatory and inhibitory activity. Computational work has shown that flatter slope is associated with an increased ratio of excitation to inhibition^43,82,83^. Further experimental work has demonstrated that the slope of hippocampal field potentials correlates to the relative density of excitatory vs. inhibitory synapses near the recording electrode^43^. Additionally, global inhibition with propofol (a GABAergic anesthetic agent) is correlated with steeping of spectral slope as measured by electrocorticography^43^.

Stroke is associated with well-characterized changes in the ratio between excitation and inhibition. During the hyper-acute phase of stroke, excitotoxicity is hypothesized to directly contribute to ischemic cell death (Reviewed by Neves et al.)^84^. In the subacute phase after stroke, animal studies have demonstrated increased tonic inhibition which likely limits recovery^85^ while potentially reducing excitotoxicity. Work in both animals^86,87^ and humans^88^ has further suggested that the longer post-stroke phase is associated with a shift toward more excitatory network behavior and that this shift is important for promoting plasticity. If indeed spectral slope reflects alterations in excitation/inhibition balance, our data would suggest that in older individuals, the chronic phase of stroke is associated with a shift toward greater inhibition (steeper aperiodic slope). It is interesting to speculate that the degree of this shift may reflect varying levels of recovery, with those having the greatest shift demonstrating the best outcomes. It is important to note however, as others have speculated^45^, that changes in slope seen after stroke may be driven by other mechanisms as well including disruption of structural connectivity between cerebral regions. Aging itself is also associated with changes in excitation/inhibition ratio. Diminished inhibitory activity has been noted with normal aging in both rodents^89^ and humans^90^. This shift may be reflected by the gradually flattening of exponent observed in our study and others^39^. Chronic stroke may negate this age-related increase in excitation/inhibition ratio as reflected by steeper spectral slope, though this clearly does not occur in an adaptive manner.

It is interesting to note that the “steepening” or increase in exponent we observed in aged individuals with stroke causes the observed slopes to resemble more closely those of the younger population (before they have undergone an age-related decline in exponent). This is not to suggest however, that stroke is in some way associated with improved cognitive performance. Indeed, a wide body of literature has shown that stroke is associated with accelerated cognitive decline^91^ as well as changes in mood^92^.

Additionally, stroke itself is an independent risk factor for dementia^8,93^. We speculate that disruption in aperiodic activity in either direction: flattening with slope with age or steepening of slope in aged stroke patients may be associated with cerebral dysfunction and that it reflects a more general disruption in neuronal activity. It is especially interesting to consider that the aged brain may be uniquely vulnerable to this kind of disruption after stroke.

Another interesting observation from our work is the relationship between lesion location and exponent. We observed the greatest effect on the exponent in ipsilesional electrodes (electrodes nearest the stroke) (Figure 8). While it is unsurprising that we observed the greatest effect of stroke nearest the stroke, it is interesting to note more widespread nature of the effect. Contralateral electrodes from the stroke group still exhibited abnormalities in the exponent, suggesting stroke mediated disruption of aperiodic network activity throughout the brain. Prior work employing both human functional imaging^28,94^ as well as animal studies^29,95^ have shown that focal stroke is associated with widespread disruption in functional network connectivity and that these disconnections may be superior predictors of deficit and recovery within certain functional domains than lesion location. Disruption of these broad networks, particularly thalamocortical networks^96^, may be a primary driver of the widespread disruption of aperiodic activity. Our observation that deep lesions had the greatest influence on cortical aperiodic activity further suggests a role for network disruption as a driver of widespread (non-focal) cortical activity changes after stroke (Figure 8). This is particularly intriguing to consider given the broad role thalamocortical connections driving feed-forward inhibition of cortical circuits^97,98^ while playing an integral role in consciousness^99^ and memory^100^. The robust influence of deep lesions on aperiodic slope we observe may represent disruption of cortical excitation/inhibition balance driven by damaged thalamocortical connectivity.

We also observed several changes within the periodic components of the spectrum in both stroke and non-stroke individuals. First, we noted fewer total peaks in the stroke group then the control group. This was most prominent when comparing young patients. Fewer identified peaks may suggest changes in spectral density are driven disproportionally by changes in aperiodic activity after stroke. We also observed a general slowing of central frequency within peaks with aging particularly in higher frequency bands. Central frequency in the alpha band was slower in individuals with stroke (Figure 5). This influence of stroke on alpha central frequency has previously been observed^45^. It is interesting that, as with aperiodic activity, we note some baseline influence of age, however, it is not as robust as the influence of stroke and age on aperiodic activity-again recapitulating prior work suggesting that aperiodic changes are a significant driver of post-stroke changes in spectral activity.

This work has several strengths including the significant number of participants, the inclusion of motor outcomes, and the wide range of ages. The study also has several important limitations. First, stroke is intrinsically, a heterogenous disease. Our data reflect a variety of stroke locations and severities. Future work will focus on further characterizing how differences in stroke location influence EEG activity.

Additionally, our data were obtained across a wide range of time points post-stroke. This limits our ability to determine what is necessarily a reflection of chronic stroke vs ongoing acute changes in stroke. Additionally, the strokes in our dataset were generally smaller. It will be important to explore these findings in larger strokes as well. Finally, future work should focus on correlating these measures to objective measures of outcomes including both motor and psycho-cognitive tests. Despite these limitations, our work builds substantially on prior work describing changes in cortical activity after stroke and characterizes a key age-dependent effect of stroke on aperiodic cortical activity. It also provides a starting point for a more granular examination of these findings in different stroke pathologies and time points as well as in non-vascular neurologic pathologies.

In conclusion, here we show that patients with stroke exhibit significant chronic changes in both the periodic and aperiodic components of the power spectrum. Our work recapitulates recent work^45^ demonstrating a shift in aperiodic activity (slope) as a significant contributor to cortical spectral changes observed after stroke. Our work adds to this work by leveraging multiple large data sets to specifically examine the influence of age on aperiodic activity. We found that the influence of stroke on exponent was primarily observed in older individuals, and that stroke had minimal effect on exponent in younger individuals. Additionally, we found the greatest influence on exponent was from deep strokes, and while the greatest effect was observed near the stroke, brain-wide changes in periodic activity were observed. Finally, our work suggests that slope in aged but not young individuals may be a relevant biomarker of motor performance.

