## Supplemental Tables for "Stroke and Motor Recovery are Associated with Regional and Age-Specific Changes in Periodic and Aperiodic Cortical Activity"

**Supplemental Table i**. Summary of the Mixed Effects Model for the Aperiodic Exponent.

```
Random effects:
 Groups   Name        Variance Std.Dev.
 subID    (Intercept) 0.063980 0.25294
 Channels (Intercept) 0.002396 0.04895
 region   (Intercept) 0.001316 0.03628
 Residual             0.018083 0.13447
Number of obs: 6519, groups:  subID, 295; Channels, 24; region, 4
---
Fixed effects:
                Estimate Std. Error         df t value Pr(>|t|)
(Intercept)     1.076751   0.029451  15.564745  36.560  < 2e-16 ***
age.c          -0.006486   0.001214 295.004984  -5.342 1.84e-07 ***
group.L         0.239294   0.029177 294.798222   8.202 7.40e-15 ***
age.c:group.L   0.003620   0.001717 295.003093   2.109   0.0358 *
---
Type III Analysis of Variance Table with Satterthwaite's method
             Sum Sq Mean Sq NumDF DenDF F value    Pr(>F)
age.c       0.51609 0.51609     1 295.0  28.540 1.838e-07 ***
group       1.21637 1.21637     1 294.8  67.265 7.402e-15 ***
age.c:group 0.08040 0.08040     1 295.0   4.446   0.03583 *

---
Signif. codes:  0 ‘***’ 0.001 ‘**’ 0.01 ‘*’ 0.05 ‘.’ 0.1 ‘ ’ 1
```

**Supplemental Table ii**. Summary of Mixed-Effects Model for Number of Narrow Band Peaks.

```
Random effects:
 Groups   Name        Variance  Std.Dev.
 subID    (Intercept) 4.919e-02 0.221797
 Channels (Intercept) 2.165e-03 0.046526
 region   (Intercept) 1.686e-05 0.004106
 Residual             7.106e-02 0.266568
Number of obs: 7080, groups:  subID, 295; Channels, 24; region, 4
---
Fixed effects:
                Estimate Std. Error         df t value Pr(>|t|)
(Intercept)     1.549357   0.020913  54.371274  74.087  < 2e-16 ***
age.c          -0.002759   0.001089 290.997258  -2.534   0.0118 *
group.L        -0.117818   0.026167 290.997262  -4.503 9.73e-06 ***
age.c:group.L   0.003668   0.001540 290.997257   2.382   0.0178 *
---
Type III Analysis of Variance Table with Satterthwaite's method
             Sum Sq Mean Sq NumDF DenDF F value    Pr(>F)
age.c       0.45646 0.45646     1   291  6.4237   0.01179 *
group       1.44054 1.44054     1   291 20.2726 9.728e-06 ***
age.c:group 0.40331 0.40331     1   291  5.6757   0.01784 *
---
Signif. codes:  0 ‘***’ 0.001 ‘**’ 0.01 ‘*’ 0.05 ‘.’ 0.1 ‘ ’ 1
```

**Supplemental Table iii**. Summary of Mixed-Effects Model for Central Frequency of Gaussian Peaks.

```
Random effects:
 Groups   Name        Variance Std.Dev.
 subID    (Intercept) 0.93596  0.9675
 Channels (Intercept) 0.01175  0.1084
 region   (Intercept) 0.04315  0.2077
 Residual             5.99837  2.4492
Number of obs: 13465, groups:  subID, 295; Channels, 24; region, 4

---
Fixed effects:
                       Estimate Std. Error        df t value Pr(>|t|)
(Intercept)           1.147e+01  1.428e-01  1.164e+01  80.299  < 2e-16 ***
band.L                7.881e+00  1.081e-01  1.336e+04  72.888  < 2e-16 ***
band.Q                2.031e+00  7.375e-02  1.343e+04  27.538  < 2e-16 ***
age.c                -1.557e-02  5.624e-03  4.545e+02  -2.767  0.00588 **
group.L               2.791e-01  1.336e-01  4.390e+02   2.090  0.03718 *
band.L:age.c         -5.562e-02  6.541e-03  1.332e+04  -8.504  < 2e-16 ***
band.Q:age.c         -7.361e-03  4.436e-03  1.338e+04  -1.659  0.09709 .
band.L:group.L        2.820e-01  1.523e-01  1.336e+04   1.851  0.06420 .
band.Q:group.L        7.056e-02  1.041e-01  1.342e+04   0.678  0.49773
age.c:group.L         3.974e-03  7.954e-03  4.545e+02   0.500  0.61761
band.L:age.c:group.L  1.061e-02  9.250e-03  1.331e+04   1.147  0.25154
band.Q:age.c:group.L  2.542e-03  6.273e-03  1.338e+04   0.405  0.68527

---
Type III Analysis of Variance Table with Satterthwaite's method
                 Sum Sq Mean Sq NumDF   DenDF   F value  Pr(>F)
band              58135 29067.7     2 13343.7 4845.9356 < 2e-16 ***
age.c                46    45.9     1   454.5    7.6590 0.00588 **
group                26    26.2     1   439.0    4.3688 0.03718 *
band:age.c          624   311.9     2 13352.9   51.9993 < 2e-16 ***
band:group           37    18.4     2 13341.5    3.0714 0.04639 *
age.c:group           1     1.5     1   454.5    0.2496 0.61761
band:age.c:group     13     6.7     2 13351.2    1.1160 0.32760
```

**Supplemental Table iv**. Summary of Mixed-Effects Model for Power at the Central Frequency of Gaussian Peaks.

```
Random effects:
 Groups   Name        Variance  Std.Dev.
 subID    (Intercept) 0.0107246 0.10356
 Channels (Intercept) 0.0001629 0.01276
 region   (Intercept) 0.0008554 0.02925
 Residual             0.0119572 0.10935
Number of obs: 13465, groups:  subID, 295; Channels, 24; region, 4

---
Fixed effects:
                       Estimate Std. Error         df t value Pr(>|t|)
(Intercept)           3.223e-01  1.727e-02  5.457e+00  18.665 3.73e-06 ***
band.L               -4.535e-02  4.902e-03  1.331e+04  -9.251  < 2e-16 ***
band.Q               -8.828e-02  3.336e-03  1.330e+04 -26.462  < 2e-16 ***
age.c                -3.989e-04  5.152e-04  3.175e+02  -0.774   0.4394
group.L               2.568e-02  1.234e-02  3.132e+02   2.081   0.0382 *
band.L:age.c         -1.304e-04  2.966e-04  1.331e+04  -0.440   0.6602
band.Q:age.c          1.791e-03  2.008e-04  1.330e+04   8.919  < 2e-16 ***
band.L:group.L        2.757e-02  6.902e-03  1.331e+04   3.995 6.50e-05 ***
band.Q:group.L       -6.577e-04  4.704e-03  1.328e+04  -0.140   0.8888
age.c:group.L        -1.310e-04  7.286e-04  3.175e+02  -0.180   0.8574
band.L:age.c:group.L  3.580e-04  4.195e-04  1.331e+04   0.853   0.3934
band.Q:age.c:group.L  3.196e-04  2.840e-04  1.330e+04   1.125   0.2604

---
Type III Analysis of Variance Table with Satterthwaite's method
                  Sum Sq Mean Sq NumDF   DenDF  F value    Pr(>F)
band             14.7156  7.3578     2 13313.8 615.3431 < 2.2e-16 ***
age.c             0.0072  0.0072     1   317.5   0.5994   0.43938
group             0.0518  0.0518     1   313.2   4.3321   0.03821 *
band:age.c        1.0992  0.5496     2 13306.4  45.9656 < 2.2e-16 ***
band:group        0.2275  0.1138     2 13306.5   9.5145 7.428e-05 ***
age.c:group       0.0004  0.0004     1   317.5   0.0323   0.85743
band:age.c:group  0.0399  0.0200     2 13306.0   1.6703   0.18823

---
Signif. codes:  0 ‘***’ 0.001 ‘**’ 0.01 ‘*’ 0.05 ‘.’ 0.1 ‘ ’ 1
```

**Supplemental Table v**. Summary of Mixed-Effects Model Exponent by Lesion Hemisphere in the Stroke Sub-Group.

```
Random effects:
 Groups   Name        Variance Std.Dev. Corr
 subID    (Intercept) 0.127184 0.35663
          contra.L    0.006810 0.08252  0.05
 Channels (Intercept) 0.003519 0.05932
 Residual             0.026972 0.16423
Number of obs: 1038, groups:  subID, 60; Channels, 20

---
Fixed effects:
                            Estimate Std. Error         df t value Pr(>|t|)
(Intercept)                1.395e+00  2.087e-01  5.573e+01   6.685 1.17e-08 ***
sex.L                      7.243e-02  7.144e-02  5.502e+01   1.014 0.315101
age                       -3.907e-03  3.372e-03  5.511e+01  -1.159 0.251469
days_to_enrollment        -3.460e-05  7.579e-05  5.503e+01  -0.456 0.649867
lesion_volume              1.336e-03  1.928e-03  5.506e+01   0.693 0.491475
contra.L                   3.770e-02  1.351e-02  7.024e+01   2.791 0.006768 **
channel_region.L          -4.940e-02  3.491e-02  1.538e+01  -1.415 0.177049
channel_region.Q           1.520e-01  3.241e-02  1.537e+01   4.689 0.000273 ***
channel_region.C          -4.509e-02  2.968e-02  1.531e+01  -1.519 0.149018
contra.L:channel_region.L  5.736e-02  1.780e-02  9.127e+02   3.222 0.001316 **
contra.L:channel_region.Q -3.511e-02  1.650e-02  9.122e+02  -2.128 0.033564 *
contra.L:channel_region.C -5.177e-03  1.499e-02  9.048e+02  -0.345 0.729854

---
Type III Analysis of Variance Table with Satterthwaite's method
                       Sum Sq Mean Sq NumDF  DenDF F value    Pr(>F)
sex                   0.02772 0.02772     1  55.02  1.0279  0.315101
age                   0.03623 0.03623     1  55.11  1.3432  0.251469
days_to_enrollment    0.00562 0.00562     1  55.03  0.2083  0.649867
lesion_volume         0.01294 0.01294     1  55.06  0.4797  0.491475
contra                0.21005 0.21005     1  70.24  7.7877  0.006768 **
channel_region        1.19858 0.39953     3  15.41 14.8124 8.308e-05 ***
contra:channel_region 0.63838 0.21279     3 911.68  7.8893 3.367e-05 ***

---
Signif. codes:  0 ‘***’ 0.001 ‘**’ 0.01 ‘*’ 0.05 ‘.’ 0.1 ‘ ’ 1
```

**Supplemental Table vi**. Summary of Mixed-Effects Model Number of Peaks by Lesion Hemisphere in the Stroke Sub-Group.

```
Random effects:
 Groups   Name        Variance  Std.Dev.
 subID    (Intercept) 0.0005591 0.02364
 Residual             0.0050348 0.07096
Number of obs: 1375, groups:  subID, 60

Fixed effects:
                                   Estimate Std. Error        df t value Pr(>|t|)    
(Intercept)                       1.454e+00  1.672e-02  6.070e+01  86.968  < 2e-16 ***
sex.L                            -9.623e-03  5.679e-03  5.655e+01  -1.695  0.09565 .
age                              -3.392e-04  2.692e-04  5.879e+01  -1.260  0.21266
days_to_enrollment               -9.512e-07  6.161e-06  6.148e+01  -0.154  0.87781
lesion_volume                     3.325e-05  1.536e-04  5.896e+01   0.217  0.82933
band.L                            1.877e-02  6.277e-03  1.360e+03   2.991  0.00283 **
band.Q                            7.710e-03  4.269e-03  1.375e+03   1.806  0.07110 .
contra.L                          1.565e-03  4.140e-03  1.340e+03   0.378  0.70549
channel_region.L                 -1.024e-04  6.241e-03  1.336e+03  -0.016  0.98691
channel_region.Q                  1.530e-03  5.861e-03  1.336e+03   0.261  0.79403
channel_region.C                 -6.333e-04  5.375e-03  1.322e+03  -0.118  0.90623
band.L:contra.L                   3.109e-03  8.309e-03  1.339e+03   0.374  0.70836
band.Q:contra.L                   2.022e-03  5.801e-03  1.335e+03   0.349  0.72745
band.L:channel_region.L          -1.514e-03  1.251e-02  1.335e+03  -0.121  0.90366
band.Q:channel_region.L           1.344e-03  8.757e-03  1.328e+03   0.153  0.87806
band.L:channel_region.Q           6.846e-03  1.178e-02  1.339e+03   0.581  0.56130
band.Q:channel_region.Q           6.751e-03  8.200e-03  1.334e+03   0.823  0.41047
band.L:channel_region.C           5.986e-04  1.084e-02  1.325e+03   0.055  0.95596
band.Q:channel_region.C           5.236e-03  7.494e-03  1.323e+03   0.699  0.48487
contra.L:channel_region.L        -1.212e-03  8.784e-03  1.327e+03  -0.138  0.89027
contra.L:channel_region.Q         7.302e-03  8.259e-03  1.334e+03   0.884  0.37679
contra.L:channel_region.C        -1.015e-02  7.622e-03  1.327e+03  -1.332  0.18307
band.L:contra.L:channel_region.L  8.794e-04  1.762e-02  1.330e+03   0.050  0.96021
band.Q:contra.L:channel_region.L -1.003e-03  1.238e-02  1.327e+03  -0.081  0.93542
band.L:contra.L:channel_region.Q  1.267e-02  1.661e-02  1.338e+03   0.763  0.44584
band.Q:contra.L:channel_region.Q  7.644e-03  1.156e-02  1.330e+03   0.661  0.50851
band.L:contra.L:channel_region.C -1.840e-02  1.535e-02  1.328e+03  -1.198  0.23098
band.Q:contra.L:channel_region.C -1.044e-02  1.062e-02  1.326e+03  -0.984  0.32539
```

```
---
Type III Analysis of Variance Table with Satterthwaite's method
                             Sum Sq  Mean Sq NumDF   DenDF F value    Pr(>F)    
sex                        0.014459 0.014459     1   56.55  2.8717   0.09565 .  
age                        0.007993 0.007993     1   58.79  1.5875   0.21266    
days_to_enrollment         0.000120 0.000120     1   61.48  0.0238   0.87781    
lesion_volume              0.000236 0.000236     1   58.96  0.0469   0.82933    
band                       0.107027 0.053514     2 1370.69 10.6287 2.627e-05 ***
contra                     0.000719 0.000719     1 1339.86  0.1429   0.70549    
channel_region             0.000676 0.000225     3 1333.32  0.0448   0.98742    
band:contra                0.002438 0.001219     2 1332.79  0.2421   0.78503    
band:channel_region        0.011957 0.001993     6 1330.49  0.3958   0.88205    
contra:channel_region      0.019837 0.006612     3 1330.75  1.3133   0.26849    
band:contra:channel_region 0.045051 0.007509     6 1328.75  1.4913   0.17747    

---
Signif. codes:  0 ‘***’ 0.001 ‘**’ 0.01 ‘*’ 0.05 ‘.’ 0.1 ‘ ’ 1
```

**Supplemental Table vii.** Summary of Mixed-Effects Model Central Frequency by Lesion Hemisphere in the Stroke Sub-Group.

Random effects:

| Groups | Name | Variance | Std.Dev. | Corr |
|---|---|---|---|---|
| subID | (Intercept) | 0.58003 | 0.7616 | |
| | contra.L | 0.03486 | 0.1867 | -0.18 |
| Residual | | 2.28228 | 1.5107 | |

Number of obs: 1453, groups: subID, 60

Fixed effects:

| | Estimate | Std. Error | df | t value | Pr(>\|t\|) | |
|---|---|---|---|---|---|---|
| (Intercept) | 1.208e+01 | 4.837e-01 | 6.065e+01 | 24.976 | < 2e-16 | *** |
| sex.L | -1.198e-01 | 1.659e-01 | 5.949e+01 | -0.722 | 0.473097 | |
| age | -1.813e-02 | 7.821e-03 | 5.980e+01 | -2.319 | 0.023847 | * |
| days_to_enrollment | 1.757e-04 | 1.782e-04 | 6.205e+01 | 0.986 | 0.327960 | |
| lesion_volume | 2.725e-03 | 4.470e-03 | 6.024e+01 | 0.610 | 0.544398 | |
| band.L | 6.924e+00 | 1.349e-01 | 1.451e+03 | 51.331 | < 2e-16 | *** |
| band.Q | 1.855e+00 | 9.116e-02 | 1.438e+03 | 20.353 | < 2e-16 | *** |
| contra.L | -1.183e-01 | 9.145e-02 | 1.822e+02 | -1.293 | 0.197550 | |
| channel_region.L | -4.734e-01 | 1.316e-01 | 1.391e+03 | -3.597 | 0.000333 | *** |
| channel_region.Q | -1.763e-01 | 1.236e-01 | 1.398e+03 | -1.426 | 0.154121 | |
| channel_region.C | -8.251e-02 | 1.135e-01 | 1.378e+03 | -0.727 | 0.467336 | |
| band.L:contra.L | -1.402e-01 | 1.782e-01 | 9.343e+02 | -0.786 | 0.431809 | |
| band.Q:contra.L | -1.840e-01 | 1.235e-01 | 1.201e+03 | -1.490 | 0.136572 | |
| band.L:channel_region.L | -8.583e-01 | 2.634e-01 | 1.397e+03 | -3.258 | 0.001147 | ** |
| band.Q:channel_region.L | -7.004e-01 | 1.845e-01 | 1.385e+03 | -3.797 | 0.000153 | *** |
| band.L:channel_region.Q | 5.033e-01 | 2.493e-01 | 1.403e+03 | 2.018 | 0.043739 | * |
| band.Q:channel_region.Q | -9.627e-02 | 1.728e-01 | 1.392e+03 | -0.557 | 0.577575 | |
| band.L:channel_region.C | -2.400e-02 | 2.292e-01 | 1.382e+03 | -0.105 | 0.916605 | |
| band.Q:channel_region.C | -1.259e-01 | 1.578e-01 | 1.375e+03 | -0.798 | 0.425210 | |
| contra.L:channel_region.L | 7.664e-02 | 1.852e-01 | 1.398e+03 | 0.414 | 0.679102 | |
| contra.L:channel_region.Q | -6.673e-02 | 1.745e-01 | 1.403e+03 | -0.382 | 0.702185 | |
| contra.L:channel_region.C | 4.063e-02 | 1.609e-01 | 1.372e+03 | 0.252 | 0.800713 | |
| band.L:contra.L:channel_region.L | 3.143e-01 | 3.715e-01 | 1.400e+03 | 0.846 | 0.397682 | |
| band.Q:contra.L:channel_region.L | 1.310e-01 | 2.608e-01 | 1.387e+03 | 0.502 | 0.615478 | |
| band.L:contra.L:channel_region.Q | 1.403e-01 | 3.513e-01 | 1.406e+03 | 0.399 | 0.689781 | |
| band.Q:contra.L:channel_region.Q | -1.814e-01 | 2.440e-01 | 1.397e+03 | -0.743 | 0.457334 | |
| band.L:contra.L:channel_region.C | -2.558e-01 | 3.247e-01 | 1.377e+03 | -0.788 | 0.431009 | |

```
band.Q:contra.L:channel_region.C  9.430e-02  2.236e-01  1.372e+03   0.422 0.673236

---
Type III Analysis of Variance Table with Satterthwaite's method
                            Sum Sq Mean Sq NumDF   DenDF   F value    Pr(>F)    
sex                            1.2     1.2     1   59.49    0.5213  0.473097    
age                           12.3    12.3     1   59.80    5.3764  0.023847 *  
days_to_enrollment             2.2     2.2     1   62.05    0.9722  0.327960    
lesion_volume                  0.8     0.8     1   60.24    0.3717  0.544398    
band                       11843.1  5921.6     2 1439.40 2594.5768 < 2.2e-16 ***
contra                         3.8     3.8     1  182.19    1.6726  0.197550    
channel_region                30.2    10.1     3 1392.62    4.4062  0.004301 ** 
band:contra                   12.0     6.0     2 1036.74    2.6285  0.072664 .  
band:channel_region          152.0    25.3     6 1386.91   11.1015 4.016e-12 ***
contra:channel_region          1.7     0.6     3 1393.16    0.2537  0.858739    
band:contra:channel_region     8.6     1.4     6 1386.70    0.6300  0.706413    

---
Signif. codes:  0 ‘***’ 0.001 ‘**’ 0.01 ‘*’ 0.05 ‘.’ 0.1 ‘ ’ 1
```

**Supplemental Table viii.** Summary of Mixed-Effects Model Power at Central Frequency by Lesion Hemisphere in the Stroke Sub-Group.

```
Random effects:
 Groups   Name        Variance  Std.Dev. Corr 
 subID    (Intercept) 0.0090781 0.09528       
          contra.L    0.0012258 0.03501  -0.07
 Channels (Intercept) 0.0002534 0.01592       
 Residual             0.0084270 0.09180       
Number of obs: 1453, groups:  subID, 60; Channels, 20

---
Fixed effects:
                                   Estimate Std. Error         df t value Pr(>|t|)    
(Intercept)                       3.904e-01  5.674e-02  5.910e+01   6.881 4.26e-09 ***
sex.L                             1.609e-02  1.946e-02  5.870e+01   0.827 0.411758    
age                              -1.192e-03  9.168e-04  5.847e+01  -1.301 0.198525    
days_to_enrollment                1.016e-05  2.072e-05  5.949e+01   0.490 0.625622    
lesion_volume                    -1.097e-03  5.241e-04  5.840e+01  -2.094 0.040602 *  
band.L                           -6.043e-02  8.423e-03  1.387e+03  -7.175 1.18e-12 ***
band.Q                           -7.855e-02  5.659e-03  1.376e+03 -13.880  < 2e-16 ***
contra.L                          1.220e-02  7.213e-03  1.248e+02   1.691 0.093333 .  
channel_region.L                  3.040e-02  1.193e-02  2.771e+01   2.548 0.016677 *  
channel_region.Q                 -5.495e-02  1.114e-02  2.840e+01  -4.932 3.23e-05 ***
channel_region.C                  8.608e-04  1.019e-02  2.822e+01   0.085 0.933255    
band.L:contra.L                  -1.263e-02  1.149e-02  1.305e+03  -1.099 0.271889    
band.Q:contra.L                  -3.897e-03  7.818e-03  1.360e+03  -0.498 0.618265    
band.L:channel_region.L           2.151e-02  1.617e-02  1.350e+03   1.330 0.183713    
band.Q:channel_region.L          -4.398e-02  1.130e-02  1.344e+03  -3.891 0.000105 ***
band.L:channel_region.Q           2.774e-02  1.534e-02  1.355e+03   1.808 0.070851 .  
band.Q:channel_region.Q          -1.342e-02  1.060e-02  1.348e+03  -1.265 0.206079    
band.L:channel_region.C           9.597e-04  1.405e-02  1.345e+03   0.068 0.945550    
band.Q:channel_region.C          -5.135e-03  9.657e-03  1.338e+03  -0.532 0.594999    
contra.L:channel_region.L         2.264e-02  1.144e-02  1.366e+03   1.979 0.048058 *  
contra.L:channel_region.Q        -8.830e-03  1.079e-02  1.367e+03  -0.818 0.413297    
contra.L:channel_region.C         1.263e-02  9.887e-03  1.344e+03   1.278 0.201615    
band.L:contra.L:channel_region.L -5.514e-02  2.289e-02  1.359e+03  -2.409 0.016135 *  
band.Q:contra.L:channel_region.L  7.460e-03  1.597e-02  1.342e+03   0.467 0.640444    
band.L:contra.L:channel_region.Q -3.756e-03  2.173e-02  1.367e+03  -0.173 0.862773    
```

```
band.Q:contra.L:channel_region.Q  2.388e-02  1.497e-02  1.348e+03   1.595 0.110904
band.L:contra.L:channel_region.C -1.136e-02  1.994e-02  1.346e+03  -0.570 0.569104
band.Q:contra.L:channel_region.C  1.304e-02  1.366e-02  1.335e+03   0.954 0.340026

---
Type III Analysis of Variance Table with Satterthwaite's method

                           Sum Sq Mean Sq NumDF   DenDF  F value    Pr(>F)
sex                        0.0058 0.00576     1   58.70   0.6834  0.411758
age                        0.0143 0.01425     1   58.47   1.6914  0.198525
days_to_enrollment         0.0020 0.00203     1   59.49   0.2405  0.625622
lesion_volume              0.0370 0.03695     1   58.40   4.3852  0.040602 *
band                       3.6879 1.84396     2 1376.62 218.8145 < 2.2e-16 ***
contra                     0.0241 0.02410     1  124.83   2.8595  0.093333 .
channel_region             0.4577 0.15255     3   27.68  18.1030  1.06e-06 ***
band:contra                0.0219 0.01096     2 1328.26   1.3010  0.272604
band:channel_region        0.1499 0.02498     6 1345.67   2.9640  0.007044 **
contra:channel_region      0.0879 0.02931     3 1364.19   3.4784  0.015459 *
band:contra:channel_region 0.0991 0.01651     6 1350.13   1.9594  0.068446 .

---
Signif. codes:  0 ‘***’ 0.001 ‘**’ 0.01 ‘*’ 0.05 ‘.’ 0.1 ‘ ’ 1
```

**Supplemental Table ix.** Summary of Mixed-Effects Model for the relationship between the aperiodic exponent and the box and block test.

Random effects:

| Groups | Name | Variance | Std.Dev. | Corr |
|---|---|---|---|---|
| subID | (Intercept) | 0.100384 | 0.31683 | |
| | contra.L | 0.003889 | 0.06236 | -0.11 |
| Channels | (Intercept) | 0.002451 | 0.04951 | |
| Residual | | 0.025505 | 0.15970 | |

Number of obs: 654, groups: subID, 37; Channels, 20

---

Fixed effects:

| | Estimate | Std. Error | df | t value | Pr(>\|t\|) | |
|---|---|---|---|---|---|---|
| (Intercept) | 9.377e-01 | 9.599e-02 | 3.859e+01 | 9.769 | 5.54e-12 | *** |
| sex.L | 5.965e-02 | 8.828e-02 | 3.701e+01 | 0.676 | 0.503407 | |
| days_to_enrollment | -9.176e-05 | 8.840e-05 | 3.697e+01 | -1.038 | 0.305974 | |
| lesion_volume | 1.287e-03 | 2.455e-03 | 3.701e+01 | 0.524 | 0.603379 | |
| bbt_affected | 1.205e-02 | 4.132e-03 | 3.728e+01 | 2.916 | 0.005972 | ** |
| age_group.L | -3.038e-01 | 1.342e-01 | 3.735e+01 | -2.264 | 0.029464 | * |
| channel_region.L | -6.180e-02 | 3.685e-02 | 3.419e+01 | -1.677 | 0.102626 | |
| channel_region.Q | 1.396e-01 | 3.419e-02 | 3.412e+01 | 4.083 | 0.000254 | *** |
| channel_region.C | -1.254e-02 | 3.124e-02 | 3.373e+01 | -0.401 | 0.690793 | |
| contra.L | 4.531e-02 | 2.386e-02 | 5.195e+01 | 1.899 | 0.063128 | . |
| bbt_affected:age_group.L | 9.883e-03 | 5.402e-03 | 3.730e+01 | 1.830 | 0.075301 | . |
| bbt_affected:channel_region.L | 6.308e-04 | 1.060e-03 | 5.648e+02 | 0.595 | 0.551958 | |
| bbt_affected:channel_region.Q | 6.564e-04 | 9.807e-04 | 5.682e+02 | 0.669 | 0.503561 | |
| bbt_affected:channel_region.C | -1.755e-03 | 8.906e-04 | 5.637e+02 | -1.971 | 0.049231 | * |
| age_group.L:channel_region.L | 1.103e-01 | 3.511e-02 | 5.645e+02 | 3.141 | 0.001771 | ** |
| age_group.L:channel_region.Q | -2.885e-02 | 3.257e-02 | 5.686e+02 | -0.886 | 0.376132 | |
| age_group.L:channel_region.C | 4.838e-02 | 2.967e-02 | 5.638e+02 | 1.631 | 0.103483 | |
| bbt_affected:contra.L | -1.261e-04 | 1.022e-03 | 5.115e+01 | -0.123 | 0.902303 | |
| age_group.L:contra.L | 2.471e-02 | 3.260e-02 | 4.881e+01 | 0.758 | 0.452216 | |
| channel_region.L:contra.L | 5.052e-02 | 3.673e-02 | 5.898e+02 | 1.376 | 0.169493 | |
| channel_region.Q:contra.L | -4.907e-03 | 3.414e-02 | 5.873e+02 | -0.144 | 0.885764 | |
| channel_region.C:contra.L | 3.108e-03 | 3.113e-02 | 5.858e+02 | 0.100 | 0.920501 | |
| bbt_affected:age_group.L:channel_region.L | -2.373e-03 | 1.500e-03 | 5.652e+02 | -1.583 | 0.114039 | |
| bbt_affected:age_group.L:channel_region.Q | 1.153e-03 | 1.387e-03 | 5.680e+02 | 0.832 | 0.405988 | |
| bbt_affected:age_group.L:channel_region.C | -2.313e-03 | 1.259e-03 | 5.635e+02 | -1.837 | 0.066708 | . |

```
bbt_affected:age_group.L:contra.L                   2.368e-04  1.409e-03  4.924e+01   0.168 0.867241
bbt_affected:channel_region.L:contra.L              8.062e-04  1.562e-03  5.882e+02   0.516 0.605887
bbt_affected:channel_region.Q:contra.L             -1.780e-03  1.453e-03  5.864e+02  -1.225 0.221201
bbt_affected:channel_region.C:contra.L             -3.617e-04  1.328e-03  5.842e+02  -0.272 0.785428
age_group.L:channel_region.L:contra.L               1.193e-02  4.962e-02  5.681e+02   0.240 0.810031
age_group.L:channel_region.Q:contra.L               1.105e-02  4.619e-02  5.686e+02   0.239 0.811047
age_group.L:channel_region.C:contra.L              -2.617e-02  4.204e-02  5.685e+02  -0.622 0.533962
bbt_affected:age_group.L:channel_region.L:contra.L  1.222e-03  2.147e-03  5.781e+02   0.569 0.569643
bbt_affected:age_group.L:channel_region.Q:contra.L -1.044e-03  1.995e-03  5.786e+02  -0.523 0.601078
bbt_affected:age_group.L:channel_region.C:contra.L  3.335e-04  1.812e-03  5.773e+02   0.184 0.854058

---
Type III Analysis of Variance Table with Satterthwaite's method
                                              Sum Sq  Mean Sq NumDF  DenDF F value    Pr(>F)    
sex                                          0.01165 0.011646     1  37.01  0.4566 0.5034074    
days_to_enrollment                           0.02748 0.027485     1  36.97  1.0776 0.3059744    
lesion_volume                                0.00700 0.007004     1  37.01  0.2746 0.6033787    
bbt_affected                                 0.21687 0.216867     1  37.28  8.5028 0.0059716 ** 
age_group                                    0.13074 0.130743     1  37.35  5.1261 0.0294641 *  
channel_region                               0.83902 0.279673     3  34.09 10.9652 3.427e-05 ***
contra                                       0.09198 0.091975     1  51.95  3.6061 0.0631276 .  
bbt_affected:age_group                       0.08538 0.085384     1  37.30  3.3477 0.0753014 .  
bbt_affected:channel_region                  0.14666 0.048887     3 565.94  1.9167 0.1257186    
age_group:channel_region                     0.49080 0.163598     3 566.18  6.4143 0.0002814 ***
bbt_affected:contra                          0.00039 0.000388     1  51.15  0.0152 0.9023026    
age_group:contra                             0.01465 0.014647     1  48.81  0.5743 0.4522162    
channel_region:contra                        0.06299 0.020995     3 588.17  0.8232 0.4813923    
bbt_affected:age_group:channel_region        0.25023 0.083410     3 566.05  3.2703 0.0209549 *  
bbt_affected:age_group:contra                0.00072 0.000720     1  49.24  0.0282 0.8672407    
bbt_affected:channel_region:contra           0.06927 0.023091     3 586.61  0.9053 0.4381860    
age_group:channel_region:contra              0.01571 0.005237     3 569.19  0.2053 0.8927067    
bbt_affected:age_group:channel_region:contra 0.02885 0.009617     3 578.08  0.3771 0.7695813    

---
Signif. codes:  0 ‘***’ 0.001 ‘**’ 0.01 ‘*’ 0.05 ‘.’ 0.1 ‘ ’ 1
```
